# A syntax network morphospace reveals sudden transitions between discrete developmental stages during language acquisition

**DOI:** 10.1101/2025.02.19.639060

**Authors:** Lluís Barceló-Coblijn, Víctor M Eguíluz, Luís F Seoane

**Affiliations:** Universitat de les Illes Balears (UIB); Laboratori d’Investigació en Complexitat i de Lingüística Experimental (LICLE-UIB); Instituto de Física Interdisciplinar y Sistemas Complejos IFISC (CSIC-UIB), Palma (Illes Balears), Spain; Departamento de Biología de Sistemas, Centro Nacional de Biotecnología (CSIC), C/ Darwin 3, 28049 Madrid, Spain; Grupo Interdisciplinar de Sistemas Complejos (GISC), Madrid, Spain

**Keywords:** language networks, syntactic complex networks, language development, Down’s syndrome, specific language impairment

## Abstract

Syntax is an aspect of human language responsible for the hierarchical ordering of linguistic structures. Syntax can be summarized by dependency trees with words as nodes and edges reflecting syntactic subordination. By merging trees from several sentences, we obtain syntax graphs or networks, which display distinct shapes depending on whether the language capability is well-formed, still developing, or pathological. Such graphs make syntactic capacity quantifiable at a systemic level, revealing emerging patterns and universalities. What is the structure of syntax networks during ontogeny in typically developing (TD) children? Do cognitively challenged children develop language through alternative routes? Here we quantify and portray the typical development of syntax networks in Dutch, and find that children affected by Down syndrome, hearing impairment, and specific language impairment initially seem to follow the typical developmental path but eventually halt, culminating in a different linguistic phenotype. Our expanded data set (with almost 50 times more data than earlier studies) and increased mathematical dimensions to quantify network shape enable us to: (i) confirm and refine a proposed sharp transition in language development, (ii) correlate specific network traits with syntax maturation, and (iii) quantify the aspects that fall short in atypical development—suggesting potential diagnostic tools. We also find grounds to hypothesize a gap in syntax maturation, separating challenged children who nevertheless reach the latest stage from others systematically stuck, regardless of their condition. Our quantitative analysis enables a rigorous visualization of linguistic development trajectories, an old (yet mostly qualitative) theme in linguistics. Similar works should allow to test and propose specific hypotheses on solid grounds, as we do here. Future efforts should generalize to other languages and/or clinical conditions, seeking patterns that might point at universalities in language development.

## I. INTRODUCTION

Language is a cognitive capability that characterizes *H. sapiens* [1, 2], carried out by dedicated circuits of both cortical and subcortical brain regions [3–10], often referred to as the *language network*. It is primarily housed in the left hemisphere, although some evidence suggests an initial potential for language to be hosted on either side [11]. Some symmetry breaking process [12, 13] must occur as the language network matures, eventually leading to salient anatomical differences between the two hemispheres. Critical periods exist during which an infant must be in active contact with language. Failing to do so may result in a seemingly lifelong inability to acquire complete language skills [14–19]. Language equips *H. sapiens* with the ability to organize thoughts, produce new messages, and convey acquired knowledge, doing so in a way that no other species can [20, 21]. Some convergences exist between mathematical properties of animal and human communication codes [22–24]. However, despite sustained training, and to the best of our understanding, other animals are unable to produce full-fledged language [25]. All this suggests that, while language is inherently human, and seemingly a biologically wired capacity, the final stages of its development require a social context and are not inescapable.

Groundbreaking research is achieving an impressive mapping of the language network within the brain— uncovering its constituent parts, what each of them does individually, and how they come together [7–10]. While generated in the brain, language is then externalized, resulting in speech, signing, or written texts. The resulting *surface form* of language presents an intricate structure that can be extremely rich and prone to ambiguity (regarding both meaning [26–28] and how words are parsed and linked together [29, 30]). Despite these intricacies, the surface form is seamlessly resolved by language users. This surface form spans several levels, from phonetics through syntax to semantics. Syntax represents the backbone that summarizes relationships, within a sentence, between words (the building blocks of language) by means of hierarchical relationships. These relationships can be captured by graphs or networks. Such graphs can be reconstructed from utterances, signings, or written text. In this framework, each lexical item is taken as a node or vertex, and a link or edge connects two nodes if a syntactic dependency relation exists between them (Fig. 1).

**FIG. 1.**
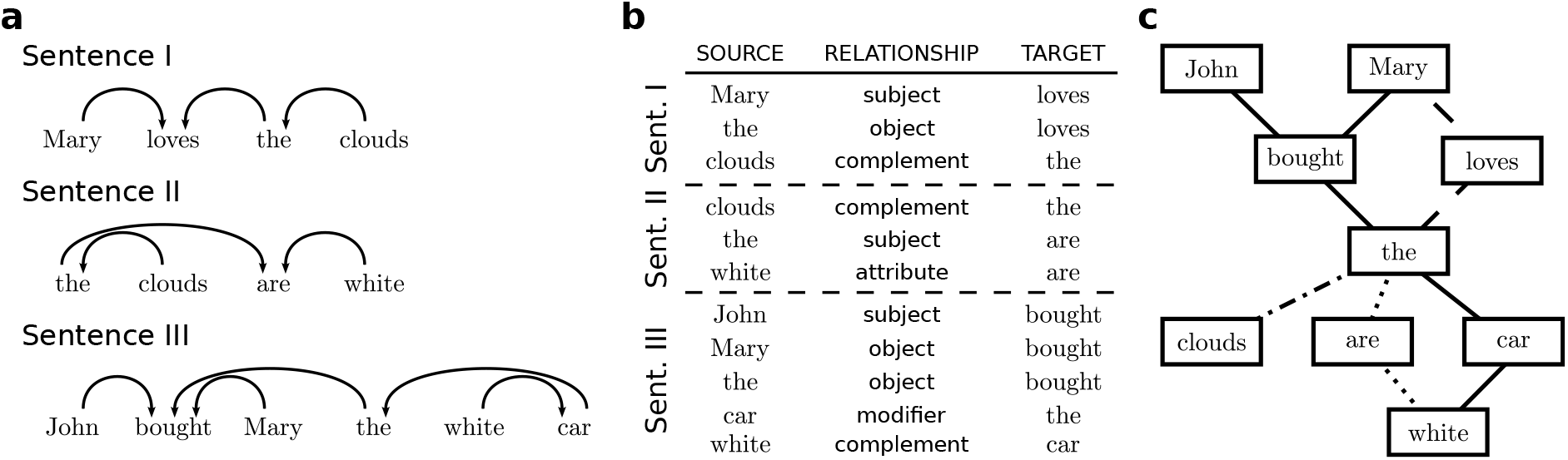
Example of syntactic analysis and reconstruction of one syntax network. **a** We start out with sentences uttered by children in a session of spontaneous conversation with their parents or caretakers. Each sentence was syntactically analyzed from within the Dependency Grammar framework. This results in a syntactic tree, which is a simple kind of graph. **b** Each word-to-word relationship was collected as the line of a plain-text file. Note that there is a directionality in this syntactic analysis. Each file contains all the data from a single conversation session, and will become a network and a data point for our analysis. **c** Each line in a file becomes an edge of the corresponding network. All words that are the same across sentences in a single session (e.g., here, “Mary”, “clouds”, “the”, etc.) are collapsed into a single node. Each two nodes are connected if there is at least one syntactic relationship between them in this session. It is still possible to reconstruct the syntactic tree of each individual sentence: they are subgraphs within the network—dashed edges for sentence I, dotted for sentence II, and solid for sentence III. Note the syntax relationship between “the” and “clouds” present in Sentences I and II, hence dash-dotted. While directionality might be kept, our current analysis did not use it.

Network theory is a very fruitful branch of mathematics [31]. A network, or graph, can recollect interactions between different components of complex systems, such as genes within a cell [32], words in a semantic space [33], species in an ecosystem [34], etc. This abstraction allows us to apply powerful computational and analytical methods derived from statistical mechanics [35], the study of graph structure [36–39], dynamical systems [40], etc.; and thus extract conclusions that might apply broadly across fields, to any networked system. Following [41, 42], here we apply such abstractions to study syntax networks generated by children as they speak.

Similarly to the brain’s language capacity, syntax networks appear to mature as children grow up [41–44], typically reaching their final form within the first four years. They start as broad trees, with large distances between arbitrary pairs of nodes. At some point, syntax networks become *small worlds* in which typical nodes are close from each other. Between these stages, the most connected words and other network properties change abruptly, similarly to a physical phase transition [41–43]. An enticing possibility is that the development of syntax networks reflects the maturation of the language organ within the brain. If this is the case, the readily accessible surface form of language could provide a window into the structures that neuroscientists are painstakingly examining. Failures in syntax formation, then, might indicate specific defects in the dedicated language circuitry in the brain. Fully achieving such a perspective remains distant for now, but it constitutes the big picture driving this work.

We examine how syntax networks mature (as abstract mathematical objects) in Typically Developing (TD) children as well as in children with developmental challenges. We provide a more detailed analysis compared to earlier studies [41–44]: We investigate additional aspects (i.e. more distinct measures) that differentiate networks from one another, and using a larger database. This allows us to tackle a series of relevant questions: What does the typical development of syntax networks (as a mathematical *graph* object) look like? Specifically so, what developmental path do they follow, from infancy to full-fledged language, across a larger, abstract space containing all possible networks?

The notion of language developing along a specific trajectory is an old theme in linguistics [45–48]. In our big picture, anatomical alterations in the language neurological structures might result in alternative linguistic development trajectories which may or may not, eventually, become capable of indistinguishable full-fledged language. We expect such alternatives to also be reflected in the developmental path of syntax networks across the mathematical space of possible graphs [49]. We lack large databases that pair the graph shape of surface syntax with the language organ structure within the brain. (At the moment, this neural network is reconstructed as averages across individuals [10].) However, atypically developing children (affected, e.g., by mutations, chromosomal trisomy, gene deletion or translocation, etc.) offer a narrow window of access. In such cases, the affected individuals may develop a visibly different phenotype—e.g. Down’s Syndrome (DS) [44]. Often, while capable of complex communication, DS children produce linguistic output that is perceived as strange or incorrect by speakers who have undergone the expected developmental steps [50].

Thus, back to questions that we can approach from our mathematical space of developing syntax graphs: Are challenged children eventually able to produce syntax networks similar to those of TD children? If not, where in the abstract space of possible graphs do atypically developed networks dwell? How far are they from typically developed ones? Is their developmental trajectory across network space similar for all children? Bringing these questions into a quantitative framework might also offer diagnostic tools.

In this paper we address all these questions from a rigorous mathematical framework. We do so by studying a large database containing language produced by: (i) 33 TD children at different stages (i.e. of different early ages); (ii) three children studied at various points in time (i.e. through longitudinal studies as they aged, producing 18, 17, and 14 data points each respectively); and (iii) three groups of atypically developing children, one containing 20 DS children, another one containing 20 children affected by Hearing Impairment (HI), and another one containing 20 children affected by Language Specific Impairment (SLI). (See App. A for details of each pathology.) From that database (introduced in Sec. II.A) we reconstructed syntax networks for each child (Sec. II.B), and performed a series of measurements that characterize the structure of each graph (Sec. II.C). In Sec. III.A we show how the abstract space of TD syntax networks is mainly shaped by a sharp transition (already identified in earlier studies [41–44]) between tree-like graphs and mature syntax networks of characteristic shape. Our abstract space allows us to visualize syntax network maturation and to compare it against graphs from atypically developing children. In Sec. III.B, we show how these atypical trajectories seem to track typical development, and stop at some point along the path, often around or right before the sharp transition. Two different interpretations are hinted at by our results: (i) that children affected by different conditions are stopped at different developmental stages (from earlier to later: DS, HI, SLI), versus (ii) that some children from either condition manage to develop syntax networks indistinguishable from TD children, and that a gap separates them from those who get halted along the way. We discuss this last possibility in Sec. III.C. In Sec. IV we contextualize our findings in the broader literature.

## II. METHODS

### A. The CHILDES data bank

The CHILDES database [51] contains numerous datasets and resources for the study of language in children of different ages, clinical conditions, and as speakers of different tongues. A prominent corpora of data created by Gerard Bol and Folkert Kuiken [52] focuses on Dutch language and includes both typically and atypically developing children. It also includes both longitudinal (which track a same child’s progress over time) and cross-sectional (which track several children of different conditions at a similar stage in time) studies. In this paper we restrict our study to a subset of the Dutch-speaking data.

Our raw data consists of transcriptions of spontaneous linguistic interactions between a child and their parent or caregiver. Each such session usually lasts between 30 and 45 minutes. Holophrases (a single-word utterance used by young children to express a complex idea or a full sentence-like meaning, relying on context and intonation to convey additional information—such as “up” to mean “pick me up”), words, or sentences were recorded. Each sentence (or equivalent unit) was stored as the line of a plain-text file. Each unique session resulted in a single file, with as many lines as sentences, words (with full sentence meaning), or holophrases were uttered. Each such single file was analyzed syntactically and processed (as described in Secs. II.B, II.C, and II.D) to produce a syntax network and a data point in our abstract mathematical space of syntax networks.

Our data comprises 141 individual sessions (which will result in as many syntax networks that will become as many data points for our analysis). Of these, 81 correspond to typically developing children, while 60 correspond to children with atypical development.

Of the 83 TD sessions, 33 of them correspond to unique sessions, each with a different child (Sup. Tab. A), while 49 of them correspond to three longitudinal studies that obtained several sessions from three different children (Sup. Tab. A). Those from unique sessions have ages between 1;07.11 (1 years, 7 months, and 11 days) and 3;04.20. For visualization purposes, we often split this data into three subgroups according to age, with 11 data points each: from 1 to 2 years, from 2 to 3 years, and from 3 to 4 years. Longitudinal studies were performed on three children labeled Daan (18 sessions, from 1;09.09 to 2;06.25), Abel (17 sessions, from 1;10.30 to 2;10.00), and Matthijs (14 sessions, from 1;11.10 to 2;05.26). Data from atypically developing children (Sup. Tab.comes from three groups of equal size (20 sessions each) of children with Down’s syndrome (from 4;04.21 to 18;11.5), children with hearing impairment (from 3;11.14 to 9;00.15), and children with language specific impairment (from 4;01.16 to 8;01.17).

We refer to App. A for table summaries of this information, as well as brief explanations of each of the atypical developmental conditions.

### B. Syntactic analysis and network reconstruction

From each of our 141 transcriptions we produced a syntax network. First, each individual sentence was analyzed adhering to the principles of Dependency Grammar [53, 54]. This procedure extracts the depedency structure beween words or morphemes with each other (Fig. 1). From each sentence, a small graph (a tree) is obtained. We collapse the trees pertaining to all sentences from an individual session by assigning each word to a node, and connecting two nodes if there is at least one syntactic dependency in one of the syntactic trees of the session between the corresponding words. The result is usually a much bigger graph than the individual trees. We term this collapsed graph a syntax network, which constitutes our unit of study.

All syntactic analyses were initially conducted manually and later validated using Netlang, a corpus annotation software [55]. Principles from syntactic theory (e.g., the Determiner Phrase [56, 57]) have been taken into account.

The resulting syntax networks were stored in plain text files either as a list of edges (with each line containing a couple of connected words, ignoring directionality) or as a list of tuples of the form (source, relationship, target) which preserved the directionality proper of syntax dependency. In any case, this directionality was ignored and we restrict our analysis to undirected networks. (Expanding it to keep directionality into account is a promising next step in the proposed analysis.) Other than that, our syntax graphs might take any form, including presenting a unique connected component, or appearing as several fragmented subgraphs.

### C. Topological measurements on syntax networks

Network science has provided us with a large toolkit to study complex networks. Many strategies are available, and our choices here do not exhaust the information that can be extracted from this data set. Our current approach is to quantify as many aspects as we could think of about a graph. Our measurements include broadly: (i) sizes, such as vocabulary (i.e. number of nodes) and edges within each net; (ii) properties pertaining to node connectivity such as their degree or clustering; (iii) properties pertaining to a relationship between nodes and the network at large, such as the graph’s diameter, assortativity, or *k*-cores; (iv) characterization of cycles present in the graph; (v) maximal constraints that can be imposed onto each network, such as the coloring number; or (vi) measurements of similarity and diversity between nodes, such as Jaccard similarities. In table I we summarize names, symbols, and literature about each of the measured properties. We provide a more thorough description of each property in App. B. We included some hints of what these abstract measurements might mean for syntax. Some of these properties might seem redundant as they are correlated—e.g., some properties increase if a graph contains longer linear chains. However, each quantity presents some potential subtlety, and all of them together are more capable of describing the different ways in which a network can be complex.

**TABLE 1.**
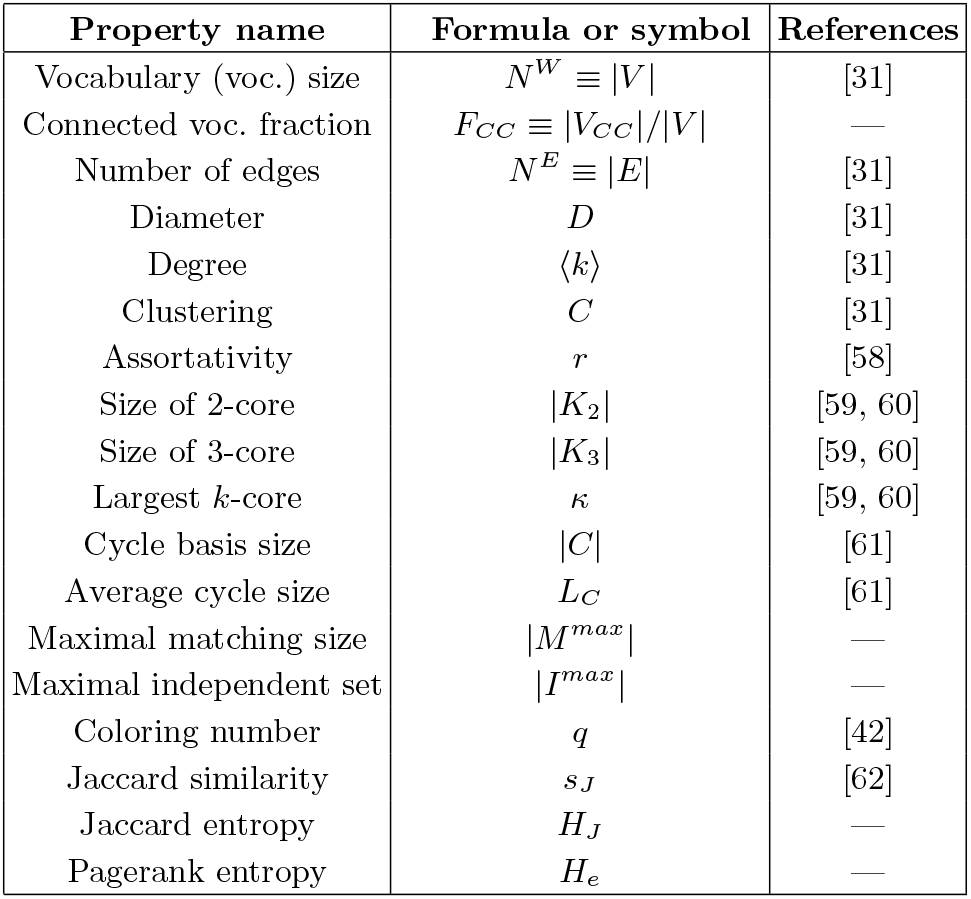
Topological properties in our analysis. Lengthier explanations with more mathematical formulations can be found in App. B.

**TABLE 2.** Fit of first PC to sigmoid Eq. 1. The mid column shows parameter estimates and uncertainties for fits to all typically development data. The right-most column shows the same for fits to longitudinal studies only.

| Parameter | All TD | Longitudinal TD |
| --- | --- | --- |
| $A$ | 10.21(80) | 15.3(2.9) |
| $B$ | -6.41(62) | -8.8(2.0) |
| $A + B$ | 3.8(1.4) | 6.5(4.9) |
| $t_{1/2}$ (days) | 795.5(9.2) | 796(17) |
| $t_{1/2}$ (years) | 2.179(25) | 2.181(47) |
| $k$ (days<sup>-1</sup>) | 0.0241(47) | 0.0152(42) |
| $Ak/4$ (days<sup>-1</sup>) | 0.061(52) | 0.058(78) |

### D. Dimensionality reduction

For each available syntax network we measured the 18 topological properties described above. These quantities characterize several ways in which graphs can be different from each other, from the obvious number of nodes (i.e. vocabulary size) and average degree (i.e. how many different syntactic relationships a word participates of), to more abstract aspects such as coloring number (which here relates to a minimal number of word classes needed to tile a network without having two words of a same class in contact) or assortativity (which gives us a hint of whether a word relates syntactically to others similar to itself).

Each graph in our data set is represented by an array containing its 18 topological measures. All networks together populate an 18-dimensional mathematical space with a cloud of points. Many of these properties are correlated—e.g. a larger number of edges likely results in a graph with more available paths, more cycles, and reduced network diameter; etc. Hence several dimensions may be redundant and the data occupies a subspace of much smaller dimension. We found a representation of reduced dimensionality, a strategy in vogue across disciplines from cell biology [63], through neuroscience [64– 66], to network topology [39]. A series of techniques such as autoencoders [67], umap [68], or non-linear manifold embeddings [69] perform this job, each with some subtleties and specific challenges. We opted for Principal Component Analysis (PCA) [70], the most straightforward and intuitive method.

In a nutshell, PCA finds the directions across the 18dimensional space across which syntax networks present more variability. Each of these directions is a linear combination of the measured properties. These directions, ranked in order of decreased variability, constitute the Principal Components (PC) of our data; and they are given by the eigenvectors of the correlation matrix between network properties, sorted in decreasing order of their eigenvalue [70]. If the data indeed occupies a smaller-dimensional subspace, then a few PC suffice to provide a satisfying description. Some bootstrapping methods are often employed to determine the optimal number of dimensions that should be kept. We forego such analysis here, as we find that a study of the first two PC allows us to tackle our research questions.

### E. Null models

We built two kinds of null models to compare our real syntax networks to, which we call *rewire* (Rwr.) and *random* (Rnd.). To build each of them, we took each real syntax network from TD children and produced 5 bootstrapped graphs.

Rwr. networks were built by randomly rewiring existing connections. These graphs were derived from uttered sentences, from which syntactic relationships were extracted. We represent connections in these networks as a list of edges given as a tuple or words (*w*_1_, *w*_2_), meaning that a syntactic dependency exists between them. To build a Rwr. network, we listed all edges, took all words in the first position and shuffled them, then took all words in the second position and shuffled them. With this bootstrapping, each word keep its original degree. Other properties might correlate with the original values as well.

Rnd. networks simply have the same average connectivity as their real-world counterpart. We took a TD syntax graph, then computed its average degree and produce 5 Rnd. graphs by generating a random network of the same size and degree. The resulting graphs might present some correlation with some properties in the original graph, but we expect them to diverge more than Rwr. from their original.

## III. RESULTS

We applied PCA to the topological properties measured on all syntax networks—including both typically and atypically developing children. This analysis gives us the main axes of variability in our data set. This means that, if atypically developing children are consistently outstanding with respect to typically developing participants, then their patterns will stand out as PC or trajectories within the resulting visualizations. Fig. 2**a** shows syntax networks from all (except two developmental studies omitted here for clarity) typically and atypically developing children projected onto our first two PC. This representation is a morphospace. Morphospaces were first introduced by Raup [71] to chart possible seasnail shells, by identifying dimensions of a 3D space with shape parameters. By moving along each dimension, shells changed in a specific way (e.g. one dimension changed length, another one increased the number of twists around the shell’s axis). This way, different morphological traits become segregated. A similar strategy has been deployed to study a host of complex systems [72–78], including language networks [28]. Here, by moving along the horizontal or vertical axes, we find differently structured networks—from simple graphs such as stars at the bottom-left (Fig. 2**b** and **c**), through expanded trees or star-like motifs connected by protruding branches around the center (Fig. 2**d-g**), to very densely connected graphs with a clear core-periphery structure towards the right half of the plane (Fig. 2**h-i**).

**FIG. 2.**
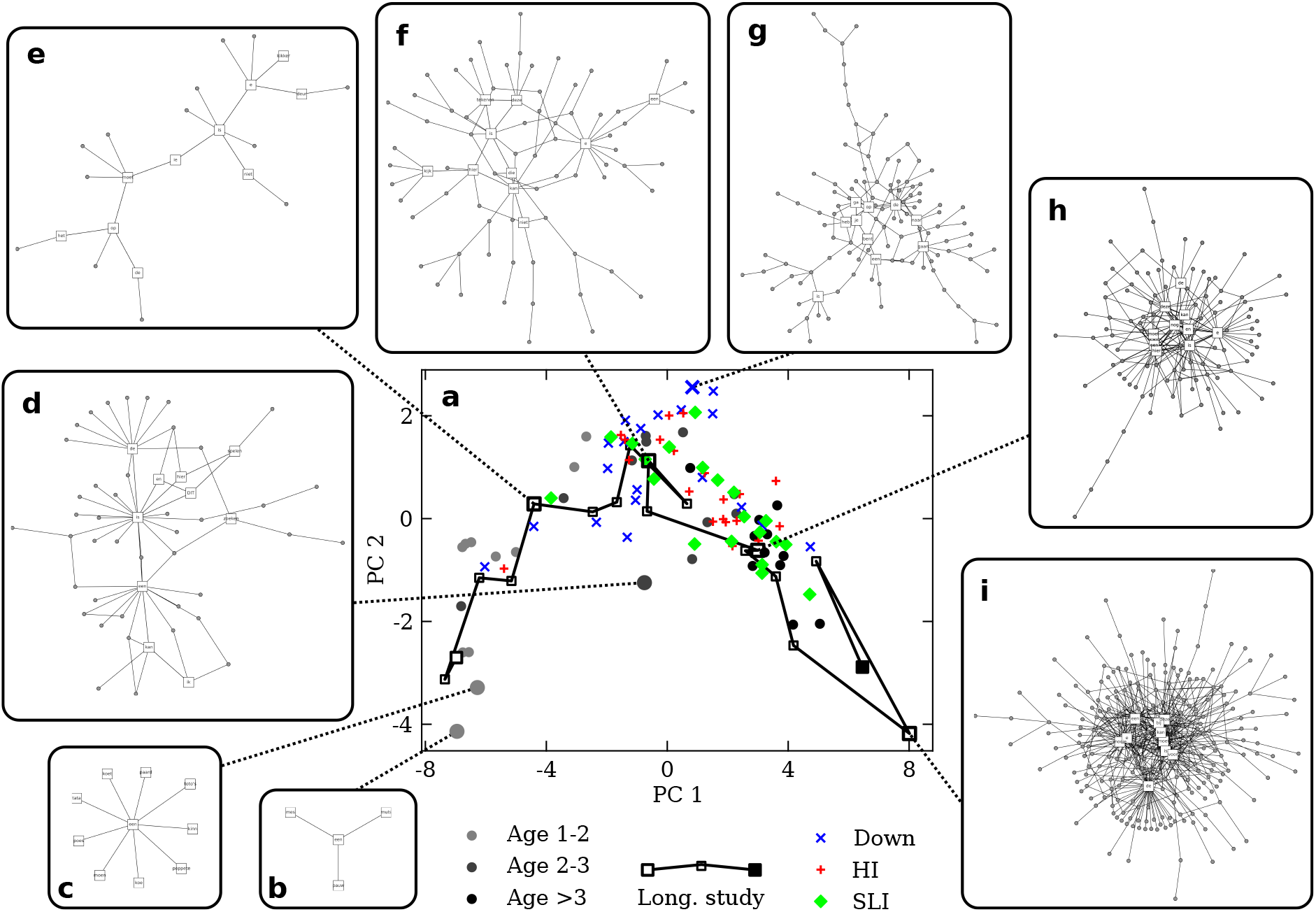
The syntax network morphospace. **a** Quantitative visualization of the syntax network development trajectory. Colors code for children of different age and condition. **b-h** Examples of real syntax graphs across the syntax network morphospace. The location of each graph in the morphospace is marked by slightly larger symbols. Boxes mark the 10 words with most connections—all other nodes appear as filled circles. **b** TD child; age 1 year, 8 months, and 23 days (∼ 628 days). **c** TD child; age 1 year, 7 months, and 11 days (∼ 586 days). **b** TD child; age 2 year, 6 months, and 10 days (∼ 920 days). **e** TD child; age 2 year, 0 months, and 4 days (∼ 734 days). **f** TD child; age 2 year, 2 months, and 16 days (∼ 806 days). **g** Atypical development (DS) child; age 18 year, 2 months, and 29 days (∼ 6659 days). **h** TD child; age 2 year, 4 months, and 0 days (∼ 850 days). **i** TD child; age 2 year, 4 months, and 0 days (∼ 905 days). Networks **e, f, h**, and **i** are from the same child.

All results in this study were elaborated including all available networks. In Sec. III.A we focus on TD children. Comparative analyses follow in Secs. III.B and III.C.

### A. A sharp transition defines the morphospace of typically developing syntactic networks

Fig. 3**a** reproduces the morphospace but including only all TD children. Any position in the plane of Figs. 2**a** and 3**a** may correspond to one (or many) graph(s) with distinct shape. However, syntax networks occupy this space irregularly, meaning that they do not present every possible graph structure—in other words, syntax networks have specific shape among all possible graphs. Our data traces an arc from left to right, with the age of children increasing throughout. This is a quantitative visualization of the trajectory of typical syntactic capacity development—i.e. of how syntax networks change shape as a child develops full-fledged language.

**FIG. 3.**
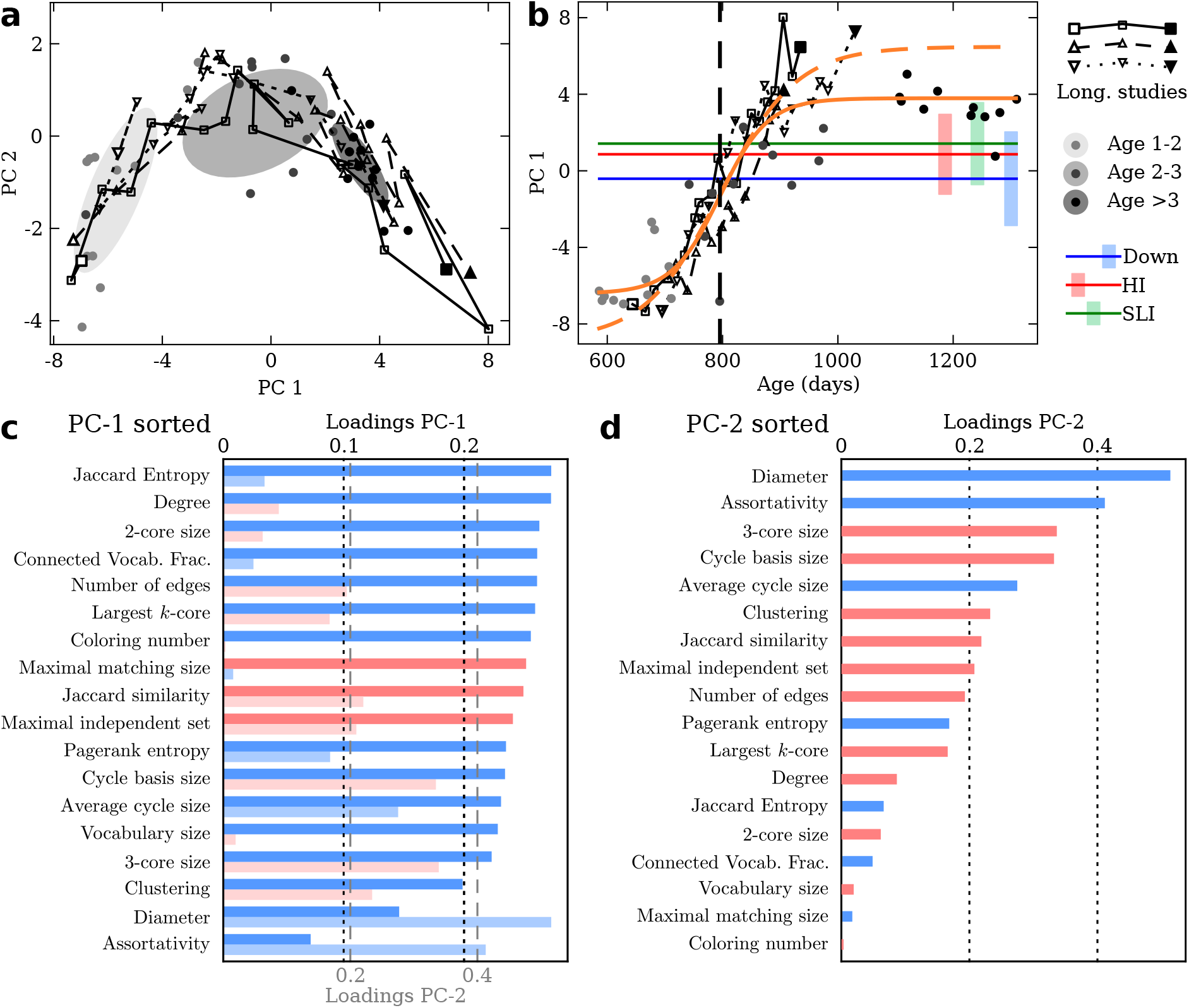
The syntax network morphospace. **a** First two PC of the syntax morphospace with graphs from typically developing children alone. Ellipsoids are centered on the mean PC-1 and PC-2 for age groups 1 ™ 2, 2 ™ 3, and*>* 3 years from lighter to darker gray. Ellipsoid axes align with, and are proportional to, the variation along the principal directions within each group. Networks from a same longitudinal study are joined by solid, dashed, or dotted curves. **b** Time evolution of the first PC for typically developing children. Horizontal lines and shaded rectangles indicate mean and standard deviation of PC-1 values in atypically developing children. Orange curves are the result of fitting sigmoids (Eq. 1) to PC-1 values of all typically developing children (solid curve) or only of longitudinal studies (dashed curve). Fit parameters with estimated uncertainties for all data can be found in Tab. II. **c** Absolute values of loadings of each property in PC-1 (darker) alongside those in PC-2 (lighter) sorted in decreasing order of PC-1 loadings. Color indicates sign: blue positive, red negative. **d** Loadings of each property in PC-2 sorted in decreasing order.

This developmental trajectory becomes more evident when PC-1 is unfolded over time. Fig. 3**b** shows that PC-1, for typically developing children, grows more or less monotonously—as we would expect from network properties that increase over time until they saturate at their *mature* value. We expect a similar behavior, e.g., for the number of words uttered within a conversation (Sup. Fig. 3**a**). Since sessions are finite, there is an average maximal number of words in any mature conversation. Sup. Figs. 3 to 9 show the temporal evolution of all properties for typically developing children. It is illuminating to see which network aspects follow the same saturating behavior as PC-1. Noteworthy among them are the fraction of connected vocabulary and number of edges (indicating that mature syntax networks reach an average maximal number of syntax relationships within finite conversations), the degree (closely related), *k*-core sizes, or the average size of cycles (suggesting that typical phrases reach a *mature* size as well).

Our PC-1 traces a much neater saturating curve than any individual property. This suggests that the main axis of variation across syntax networks captures the maturation process. Fluctuations (expected and observed in individual properties) are smoothed out in PC-1. The resulting trajectory is relatively sharp as well. Orange curves in Fig. 3**b** are fits of sigmoids with the form

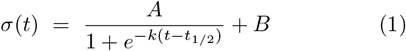

to PC-1 values from all data in TD children (solid) and to longitudinal cases only (dashed). *A* + *B* represents the saturating value of PC-1; *t*_1*/*2_ is the time at which the maturation process reaches its half, which is also when development happens at its fastest rate; and *Ak/*4 is the slope of the curve at *t*_1*/*2_ (i.e. the maximum rate of change). Tab. II shows fit results for all parameters with uncertainties.

Such a sharp development has been compared to a *phase transition* in the topology of maturing syntax networks [41, 42]. Our work corroborates this result: language development is sudden in infants, and saturates to a completion—which should be similar to its adult form. Our inclusion of more topological properties allows us to obtain better defined transition curves than before, and thus to better estimate the transition point, *t*_1*/*2_. We locate it at ≃ 2.179 ± 0.025 years for all data and at ≃ 2.181 ± 0.047 years for longitudinal studies only. These values are quite similar and within uncertainty ranges, even though children from longitudinal studies saturate to higher values of PC-1. This higher saturation value might relate to increased engagement, since these children were repeatedly interviewed under similar circumstances. It seems relevant that the transition point, *t*_1*/*2_, remains so similar. The maximal development rate, *Ak/*4, is also quite close; at ± 0.061 0.052 and 0.058 ± 0.078 respectively, and within uncertainty ranges. This supports the presence of a cognitive phasetransition-like behavior that is the same for all children— despite possible differences in engagement.

Fig. 3**c** shows the loadings of PC-1. Loadings tell us how much each original measured property (Tab. I) correlates with PC-1, which seems to capture development. We find that almost every variable has very large loadings, meaning that most network traits follow a monotonic trend towards their mature value. Three of these properties (maximal matching size, Jaccard similarity, and maximal independent set) decrease monotonically, meaning that their values decrease as syntax graphs mature. For Jaccard similarity, this means that, in early syntax networks, words have more similar neighborhoods than later.

While all topological traits are rather monotonic, PC-2 is not—neither with respect to PC-1 (Fig. 3**a**) nor to time (Sup. Fig. 2**b**). This means that, while the development process (and, specifically, the maturation of most topological properties separately) is rather straightforward, there is a combination of factors in the data that starts with some low value, peaks during maturation, and then wanes. This raise-and-fall signature is the second most important feature of the data (since it is captured by PC2), but it could easily be missed if we would look at any individual property alone.

If we think about the maturation process portrayed by a sigmoid in PC-1, such raise-and-fall behavior is displayed precisely by its derivative: it is small for *t* ≪ *t*_1*/*2_ and *t* ≫ *t*_1*/*2_ and it peaks at *t*_1*/*2_. It is an enticing possibility that PC-1 shows the level of syntax maturation while PC-2 shows ratio of development. However, we have not been able to establish this with our data. A limitation to do so is that PC-2 over time is much noisier than PC-1 (Sup. Fig. 2**b**).

Fig. 3**d** shows the loadings of PC-2. This component is made up mainly by the two properties least correlated with PC-1 (diameter and assortativity) plus combinations of all other monotonic properties with changing signs (which is needed to craft something that rises and falls out of similar monotonic trends).

Network diameter, the property with largest loading on PC-2, is particularly interesting (Sup. Fig. 5**a**). It starts out necessarily small, as syntax networks begin with few words. It grows monotonically as children utter more sentences. But then it stops growing and even decreases slightly (which is difficult to appreciate in Sup. Fig. 5**a**). This behavior becomes more clear if we normalize by the number of nodes in the largest connected component— i.e. if we present the diameter as a ratio to the number of connected vocabulary (Sup. Fig. 10). We see that this number drops drastically after the transition point. This means that, beyond this point, syntactic networks are able to grow without increasing their diameter. In other words, while vocabulary grows, the graph remains well connected and accessible.

**FIG. 4.**
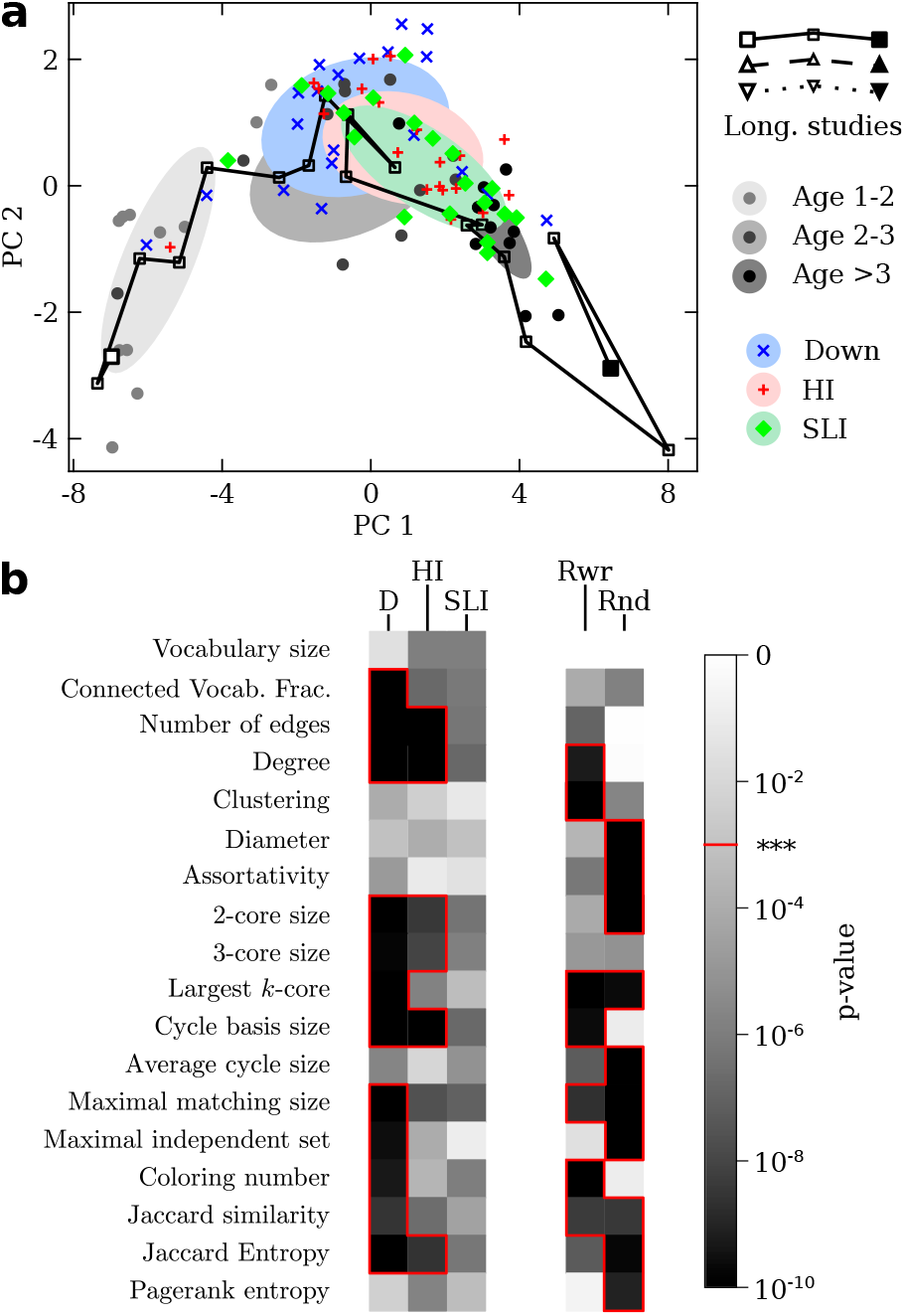
Analysis of atypically developing children. **a** Sames as in 4**a**, but including data from atypically developing children. The added ellipsoids are centered on the mean PC1 and PC-2 for children affected by Down syndrome (blue), hearing impairment (red), and specific language impairment (green). Their axes align with, and are proportional to, the variation along the principal directions within each group. **b** Reports of*p*-values from a*t*-test comparing each graph property from each atypically developing group to the same property in the group of oldest, typically developing children (ages 3 ™ 4). Properties with a *p*-value below 10^−3^ are enclosed in red circuits. Same test are reported for null-model networks.

### B. Atypical development tracks, and stops along, the typical development trajectory

A possibility with atypically developing children was that they might follow alternative maturation paths, and that they might end up in very different corners of the morphospace. This does not happen. Fig. 4**a** shows the morphospace with all syntax graphs, including those from atypically developing children (two developmental studies are omitted again for clarity). Data from atypically developing children is distributed close to and alongside the TD trajectory. This suggests that they retrace the expected (typical) developmental path. But the final end of the arc is not too well populated, hinting that atypically developing children might cease maturing at different points along the way. Ages of HI and SLI children are between 5 and 10 years, while most DS children are well in their teens (see Sup. Tab. A). Compared to typically developing children, they had more time to reach their final linguistic stages—but no guarantee exists that they did.

Colored ellipsoids in Fig. 4**a** are centered at the mean PC-1 and PC-2 values of each group. According to them, DS children (blue) halt their progress earlier than HI children (red), who in turn stop slightly earlier than SLI children (green).

Fig. 4**b** compares each graph property in atypically developing children to the same property in the TD group with oldest children (ages 3 to 4). The gray scale shows *p*-values of a double-tailed *t*-test from each comparison, with *p <* 10^−3^ marked in red. The smaller a *p*-value, the less likely that the differences between atypical and typical syntax networks is due to chance. We need to be careful with analyses based on *p*-values, but for economy of language we will say that whenever *p <* 10^−3^ then two properties are distinguishable between typical and atypical children. If *p >* 10^−3^ we will say that that property is indistinguishable or similar in both groups, and that differences are likely due to random fluctuations.

An outstanding observation is that the SLI group presents a *p*-value larger than 10^−3^ for all properties. This indicates that this group is more similar to children with typical development, and that differences between SLI children and those aged 3 ™ 4 are perhaps due to chance. In turn, HI children present certain properties with *p*-values below 10^−3^, and DS children present those same properties and a few additional ones. This recovers the order in which each group halted along the development trajectory. It is informative to look at what properties differ one by one.

All children reach similar vocabulary sizes, but only DS speakers fail to integrate a similar fraction in the largest connected component. This suggests that their linguistic output contains more disconnected structures. Both DS children and HI children have fewer edges and a lower average degree, indicating that they form fewer syntactic connections than other children.

2and 3-cores are typically smaller in both DS and HI children (even though HI children have similar largest *k*-cores, *κ*, than TD children). This indicates that DS and HI children often fail to establish large dense cores in their syntactic networks, and that their networks contain more terminal branches (which are linear chains that disappear easily in the process of finding *k*-cores). This is consistent with the reduced size in cycles basis also observed for both DS and HI children. Meanwhile, the average cycle length is indistinguishable—i.e. they close less cycles but those closed have similar lengths as in others.

Only DS children have different constraint properties from others. Specifically, they have larger maximal matching and independent set sizes and smaller coloring number. This is again compatible with a network with terminal branches or linear chains.

Interestingly, DS children also have a higher Jaccard similarity than others, suggesting that their words are less different from each other. Finally, again both DS and HI children present a lower Jaccard entropy.

We generated null models (Sec. II.E) by shuffling all the connections in a given real graph (Rwr.), or by producing graphs that had the same average connectivity than real syntax networks (Rnd.). We can think of null models as specific kinds of pathological syntax networks. It is then interesting to compare which properties differ with respect to those of mature syntax graphs (rightmost columns, Fig. 4**b**). These *p*-values indicate that null models are different to mature syntax networks in distinct ways than networks from atypically developing children, showing that the morphospace of possible graphs is much larger than that explored by syntax.

### C. Discrete developmental stages

The distribution of data points seems irregular along the syntax development trajectory (Fig. 5**a**). They are scarce in the first third of the arc, but denser both near its peak and at a point along its final third. This aligns well with both classic [45, 79] and network-based [41, 43, 44] studies on language acquisition that strongly suggest discrete developmental stages. We compare our findings to these previous studies in Sec. IV.

Trying to recover such discrete developmental stages, we fitted a *k*-means model (with *k* = 3) to our data. Fig. 5**b** shows results in which the model was fitted to all TD children. While only our main two PC are plotted, we used all components to fit the model. We also tried out different variations—specifically: fitting only to PC-1 and PC-2, fitting to all typically and atypically developing children, fitting to atypically developing children only; and combinations thereof. Our results were consistent in all cases.

Our 3-means (orange circles in Fig. 5**a-b**) identifies the scarcely sampled first third of the development trajectory (Stage I), the densely populated region near the trajectory’s peak (Stage II), and the other densely populated region towards the end of the trajectory (Stage III). Stage II corresponds to a point right before the moment of fastest development (*t*_1*/*2_). Stage III corresponds to a point on the fully-matured side of the trajectory.

While these three stages were derived from typically developing children, atypically developing participants fall seamlessly into the two final phases (save one DS and one HI children stuck in the first one). They show the same accumulation points right around the peak of the trajectory and after it. Children at each stage become indistinguishable (in a statistical sense) from TD children at the same phase. This is so for children of any condition (DS, HI, and SLI).

**FIG. 5.**
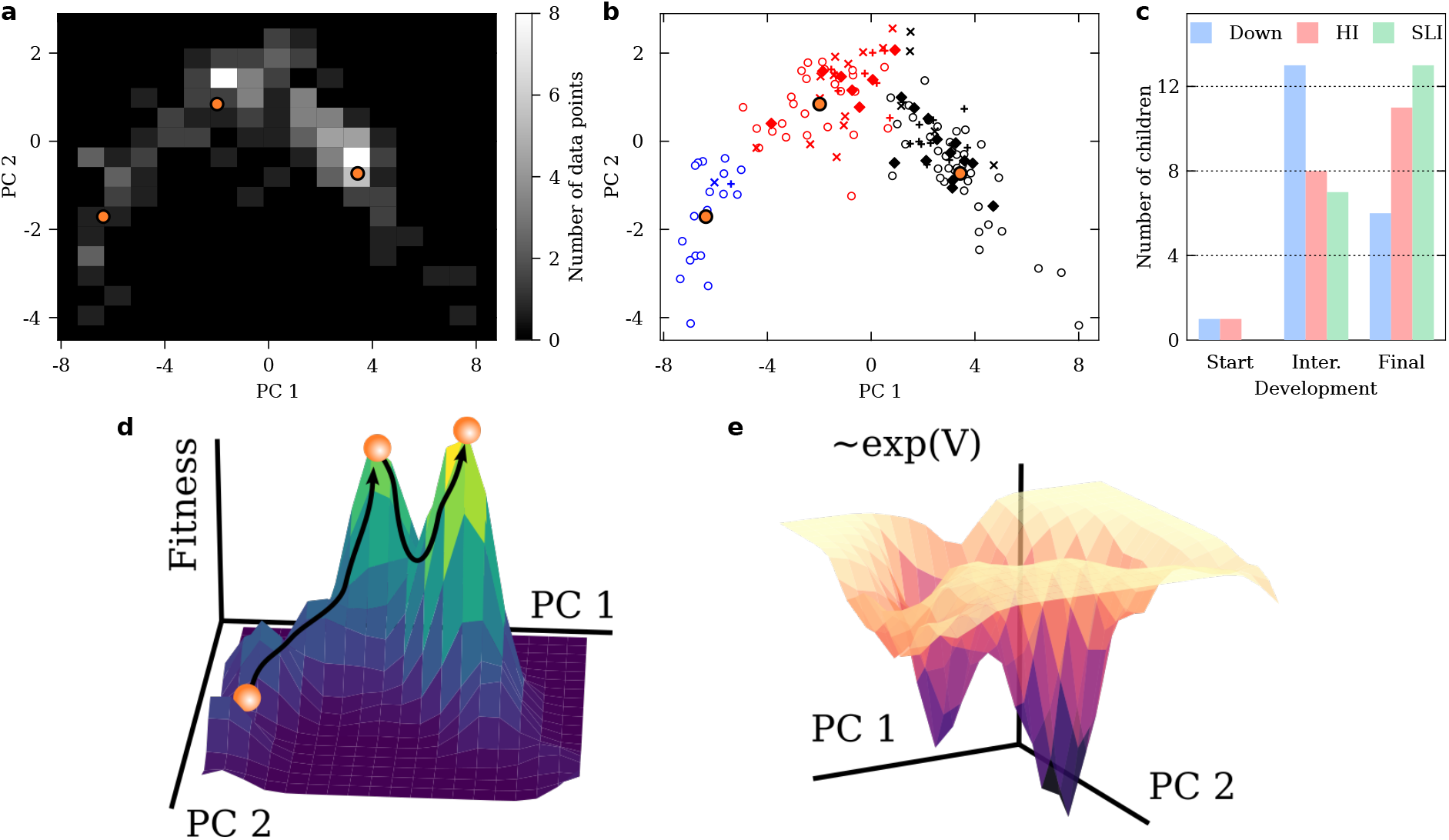
Identifying discrete developmental stages. **a** An histogram with the number of graphs found along the developmental trajectory suggests that discrete maturation stages exist. Orange circles indicate the centroids of a 3-means model fitted to the data. **b** All data points classified after the developmental stages derived from the 3-means model. Empty circles are typically developing children. Atypically developing children are shown with their original markers: crosses (DS), plus signs (HI), and diamonds (SLI). **c** Number of atypically developing children in each 3-means-derived stage. **d-** Approximation of a fitness landscape (**d**) and an energy potential (**e**) from our empirical data. These plots are for illustrative purposes only—they are not mathematically rigorous, but they convey these concepts and are ellaborated with the actual empirical data. The fitness landscape consists of the histogram in panel **a** smoothed with a Gaussian filter of unit standard deviation in every direction. Balls rolling over the fitness landscape represent a made-up child following a developmental trajectory. The energy potential (rather, its exponential) is the negative of the fitness landscape.

This suggests an alternative, very appealing reinterpretation of our data. In the previous section we assumed a smooth development along a continuous trajectory. But phase transitions are discreet. Appearances of continuity come from noise or flimsy transitory states caught as they are traversed irreversibly. Fig. 5**b** suggests that children of any condition (typically or atypically developing, and among them those with either impairment) can instead be grouped according to whether they managed to reach the full-fledged stage or not, with some obvious noise and variation.

According to this interpretation, DS, HI, and SLI children would not halt collectively at different lengths along the development trajectory. Rather, individual children of each condition have a different probability of bridging from Stage II to Stage III. Fig. 5**c** shows the number of children with each condition at each stage. It seems more difficult for DS children to make the final transition, while this is easier for HI and SLI children. Since there are more DS children stuck in the second stage, when averaging over all DS participants it seems that they, as a whole, halt earlier along the development process.

At the moment, our data is suggestive of these two different interpretations. However, the boundaries between the second and third stages in Fig. 5**b** are blurred. This is likely because of some of the mechanisms suggested above that can make discontinuous phase transitions look smooth (i.e. noise or transients caught in the act). Our second interpretation, which proposes three discrete developmental stages, is much more appealing. It also aligns better with the existing literature on language acquisition and corroborates the findings of other studies on network science applied to syntax development.

## IV. DISCUSSION

We have analyzed the largest database to date to quantitatively chart the developmental trajectory of human syntax networks. Our study produced a morphospace of syntax graphs—i.e., a mathematical space in which the structure of different networks is mapped onto points in a plane, and in which moving along specific directions we observe how certain traits of the structure of graphs vary (Fig. 2**a**). For example, moving from left to right changes: (i) from smaller to bigger networks; (ii) from nodes (words) that have less neighbors in average to others better connected, which are also part of denser cores; (iii) from graphs that present less and shorter cycles to others with more and longer loops; etc. Moving from bottom to top instead shifts from (i) networks with smaller diameters (whether they have less nodes, at the left of the plane; or more, but are tightly connected, at the right) to others with larger diameter; (ii) from graphs which are more assortative to others that are more disassortative; etc.

Each point of this morphospace corresponds to networks with distinct shapes. In principle, we could have as much graph diversity as room there is in the plane. But real syntax networks from both typically and atypically developing children do not occupy the morphospace uniformly. They trace an arc that raises and falls. The age of TD children increases along this arc—suggesting that the main dimension of the morphospace, PC-1, tracks development. Indeed, PC-1 over time unfolds like a sigmoid (Fig. 3**b**), with a sharp increase at some time, *t*_1*/*2_, and a saturation point reached for *t > t*_1*/*2_. This is what is expected of a trait that matures to completion over time. The sharpness of the sigmoid is consistent with a phase transition in the topology of syntax networks, as identified in earlier works [41–44]. We put forward the hypothesis that, while PC-1 traces maturation, PC-2 might be tracing development rate. However, noise in PC-2 over time prevents us from establishing this on a sound basis.

Additionally to TD children we studied 20 children affected by Down’s syndrome, 20 affected by hearing impairment, and 20 affected by language specific impairment. People affected by these conditions (including adults) often produce linguistic output that is identified as odd by TD speakers. A possibility would be that they produce radically different syntax networks, perhaps inhabiting remote corners of the morphospace. This does not happen. All non-typical groups appear to fall close to the TD developmental trajectory, and they seem to halt their development at different lengths along this process. The order in which each group halts depends on the clinical condition: the DS group stops earlier than HI, which in turn stops earlier than SLI.

An enticing possibility (taken here as a working framework) is that syntax network development might reflect maturation of the language organ within the brain— hence that our syntax graphs serve as proxies for syntactic endophenotypes [49]. Then, the length along development at which different children stop should correlate with differences in brain structure (which exist, as evidenced by brain studies on clinical populations [80, 81]). Studies pairing surface syntax of individuals with neuroimaging of their language organ (currently a technical challenge) are needed.

We looked at which network properties are different in atypically developing children with respect to our oldest TD group. These differences might serve as diagnostic criteria. They reflect that syntax network shape varies— e.g., between DS and full-fledged TD language. These differences, in turn, must stem from the distinct ways in which each child builds sentences. DS children use some mechanisms that are less common in TD children [44]. For example, they abuse coordination; while typical syntactic hubs (such as determiners or pronouns) often fail to establish enough connections in DS children. This loss of hubs might result in more fragmented graphs, which could explain why DS children have less vocabulary in their largest connected components, less average edges, and lower degree; among other differences. On the other hand, the abuse of coordination (think, long sentences connected by “and”) might result in lengthy terminal branches protruding out of a network core, which could explain discrepancies such as maximal matching and independent set sizes. We are unaware of coping mechanisms in HI or SLI children, but they should be considered when trying to explain the observed differences.

An alternative interpretation of our results suggests three distinct developmental Stages (labeled I, II, and III). From this perspective, atypically developing children do not appear to halt their progress at gradual lengths along the trajectory. Instead, each specific child either gets stuck in Stage II or makes the jump to Stage III (were most TD children with full-fledged language dwell). When examining graph traits (e.g., degree, clustering, etc.), children at each Stage present very similar networks, regardless of their impairment. For example, DS children in Stage III exhibit syntax graphs indistinguishable (in the statistical sense mentioned in Sec. III.B) from TD full-fledged language. (We refer to indistinguishability of graph properties—it remains possible that Stage III DS children are still given away by nuances that do not alter syntax noticeably).

These discrete development phases are very evident in our data (Fig. 5**d-e**), but some noise and intermediate cases are present. Further study is necessary, because the existence of discrete Stages in language development touches upon hotly debated issues in the study of language acquisition, as well as in contradicting views of genetic determinism versus Evo-Devo and phenotypic plasticity [82, 83].

A classic view of language acquisition, championed by Tomasello [84, 85], posits that the process is “lineal”, that there are no discrete jumps as language moves towards its full-fledged state. On the other hand, both classic [45, 79] and network-based [41–44] approaches seem to identify several different stages. In a classic work, McNeill [79] identifies several stages of language development before full maturation, among them: one consisting of holophrases (see Sec. II.A) likely undetected in our data; another displaying two-word combinations between and 2 years (akin to our Stage I) and a more complex phase with chains of more than 2 lexical items between 2 and 3 years (which may fit our Stage II). Two earlier network-based studies [41, 42] identify a sudden regime shift, which is compared to phase transitions from physics (and termed a “chromatic transition” in [42], where the coloring of the graph, which we also study, is identified as a relevant property). The interpretation that better fits our observations is that these works subsumed the progression between Stages I and II before the transition, and that this transition captures a sudden shift between Stages II and III. Other network-based studies by one of us (LB-C, et al.) already hinted at three linguistic phases [43, 44], albeit with less conclusive data and based on less network properties. Specifically, it was suggested a first phase in which words in the largest connected component had on average one edge, a second one with more than one edge but less than two, and a third one with an average degree of 2 or more. We speculate that our current analysis renders a more clear picture of those same phases, hence accumulating evidence in favor of discrete progression of language acquisition. Tellingly, the studies in [43, 44] involved more linguistic variation, suggesting a language-independent developmental pattern.

A useful tool for conceptualizing phase transitions are energy potentials and their reverse, fitness landscapes. We informally derived such objects from our data as an illustration (Fig. 5**d-e**). In a fitness landscape (Fig. 5**d**), graphs that implement their function better have a higher fitness. A developmental trajectory is conceptualized as a ball rolling uphill. In an energy potential (Fig. 5**e**) the same trajectories are given by balls rolling downhill—both representations are conceptually equivalent, but biologists often prefer the former and physicists the later. In either representations, the two last developmental stages appear as prominent peaks, suggesting some sort of “cognitive attractors”—i.e. ways of operating, cognitively, that fulfill some certain function perhaps somehow optimally. From this perspective, escaping the attractors in Fig. 5**d-e** would be difficult due to valleys or barriers that need to be traversed from one working configuration to the other.

Adopting a modern Evo-Devo perspective, with the hope of igniting debate, we launch two bold conjectures that should be put to further test.

First, similarly to how embryo development often retraces earlier evolutionary steps, we suggest that language organs might also revisit previous evolutionary linguistic phases as they mature. If so, then the “cognitive attractor” of Stage II could resemble an earlier phase in language evolution [86], more primitive than our present full-fledged capacity. The debated possibility of a protolanguage comes to mind [87, 88]. As already suggested by Bickerton in [87], the study of infant language would be an opportunity to understand the early evolution of the language capacity.

Second, we bring up Pere Alberch’s ‘logic of monsters’ [89, 90], which posits constraints in evolution and development that surface particularly in pathological phenotypes. We find very compelling that atypically developing children that do not reach Stage III seem captured by the attractor around Stage II *irrespective of their condition*. This suggests an intrinsic logic to both early and faulty syntax that might be difficult to escape. The statistical indistinguishably between syntax networks of Stage II TD infants and Stage II DS, HI, or LI children suggests that some strong constraints (perhaps rooted in the mathematics of network theory) are at play.

Due to space constraints and clarity, we focused on the analysis of the first two PC. But the data set and methodology presented here is not exhausted by our current work. On the one hand, we could explore differences in the next PC, but their importance dwindles with rank (PC-3 is a less prominent feature than PC-2, and so on). The pioneer work on the evolution of syntax graphs [41] included a study of how certain words make their way through the network to become hubs. We have not looked at individual words here—again because of space and time constraints. Such an analysis should be carried out in the future, and it should cast some light on the different specific syntactic mechanisms along the development trajectory. Also, this study deals with Dutch-speaking children. We assume that there is nothing particular about this tongue, and that our results should hold for any monolingual group of children acquiring language. However, similar studies should be implemented to confirm our findings or to find interlinguistic differences.

## Acknowledgments

LB-C was supported by grant PID2021-128404NA-I00 funded by MCIN/AEI/10.13039/501100011033. LFS received funding from the Occident Foundation (grant FJSCNB-2022-12-B), and from the Spanish State Research Agency, AEI, through the “Severo Ochoa” Programme for Centres of Excellence in R&D (CEX2023-001386-S) and, together with the Spanish Department for Science and Innovation (MICINN), through project PID2023-153225NA-I00 within the State Program for Knowledge Generation.

## Appendix A: Details of participants and atypical development conditions

Down’s syndrome occurs in 1 in 700 live births. It is a syndromic condition of known etiology. The most typical genotype of DS carries an extra copy of chromosome 21 (Trisomy 21). Other profiles, such as translocation and mosaicism, might result in DS but are less common. It has been extensively studied how DS influences language development [52, 91–93]. Several studies point to a less affected vocabulary and a more impaired syntactic capability [94].

Language development in children affected by hearing impairment is inevitably affected, since the input produced by others is not processed under the same conditions as in TD children [52, 95]. Speakers with HI have been observed to experience difficulties in processing both sentential syntax and morphosyntactic structures [95, 96].

Specific Language Impairment [97] (also known as Developmental Language Disorder) has been extensively studied since the 1980s and is one of the most researched clinical conditions in the field of clinical linguistics. SLI is a syndrome observed in children who exhibit atypical ontogenetic linguistic development without any apparent non-linguistic causes. These causes could include, for example, neurological dysfunction, general mental or cognitive delay, hearing impairment, or inadequate or insufficient exposure to linguistic stimuli due to the socio-educational characteristics of the environment in which SLI children are raised [98].

## Appendix B: Detailed description of topological properties

Let us introduce some useful notation: A graph or network, 𝒢, consists of a set, *V*, containing *N*^*n*^ ≡ *V* nodes or vertices *v*_*i*_ ∈ |*V* | ; and a set, *E*, containing *N*^*e*^ ≡ | *E* | unordered tuples of the kind (*v*_*i*_, *v*_*j*_) ∈ *E* that indicate that nodes *v*_*i*_ and *v*_*j*_ are connected. We call the elements of *E* edges or links indistinctly. We will say (*v*_*i*_, *v*_*j*_) ∈ *E* if either (*v*_*i*_, *v*_*j*_) ∈ *E* or (*v*_*j*_, *v*_*i*_) ∈ *E*. For simplicity, we might note nodes *v*_*i*_ and *v*_*j*_ as just *i* and *j* at odd times—this notation will be clear from the context. It is convenient to introduce the adjacency matrix: *A* ≡ {*a*_*ij*_; *i* = 1, …, *N*^*n*^; *j* = 1, …, *N*^*n*^} with *a*_*ij*_ = 1 if (*v*_*i*_, *v*_*j*_) ∈ *E*. It is also convenient to introduce the neighborhood of a node: *V*_*i*_ ≡ {*v*_*j*_ ∈ *V*, (*v*_*i*_, *v*_*j*_) ∈ *E*}. In this paper we only work with unweighted, undirected graphs with no self-loops (hence (*v*_*i*_, *v*_*i*_) ∉ *E* for any *i*).

**TABLE I.** Summary of TD participants, individual sessions. Data collected at 1 (left), 2 (center), and 3 (right) years of age. Age format is year;month.day. Participants are sorted by increasing age.

| 1-2 years | 2-3 years | 3-4 years |
| --- | --- | --- |
| Barry 01;07.11 | Dieuwertje 02;00.12 | Bert 03;00.12 |
| Tijn 01;07.16 | Katja 02;01.09 | Katja 03;00.15 |
| Marco 01;07.20 | Marco 02;01.22 | Maaïke 03;00.25 |
| Corine 01;08.06 | Tijn 02;02.05 | Floor 03;01.24 |
| Gea 01;08.23 | Channe 02;02.29 | Frederique 03;02.18 |
| Thom 01;10.02 | Karel 02;03.16 | Marcel 03;04.16 |
| Ronnie 01;10.05 | Rianne 02;04.20 | Otje 03;04.20 |
| Els 01;10.12 | Els 02;05.07 | Chantal 03;05.10 |
| Suzanne 01;10.17 | Maaïke 02;06.10 | Jelle 03;05.28 |
| Rogier 01;11.16 | Floor 02;07.27 | Martijn 03;06.06 |
| Marieke 01;11.12 | Frederique 02;08.05 | Bert 03;07.05 |

**TABLE II.** Summary of TD participants, longitudinal studies. Data for Daan (left, 18 sessions), Abel (center, 17 sessions), and Matthijs (right, 14 sessions) over multiple observations. Age format is year;month.day. Participants are sorted by decresing number of sessions.

| Daan | Abel | Matthijs |
| --- | --- | --- |
| 01;09.09 | 01;10.30 | 01;11.10 |
| 01;10.01 | 01;11.26 | 01;11.24 |
| 01;10.16 | 02;00.11 | 02;00.09 |
| 01;11.21 | 02;01.02 | 02;00.24 |
| 02;00.04 | 02;01.16 | 02;01.07 |
| 02;00.22 | 02;02.19 | 02;01.21 |
| 02;00.29 | 02;03.02 | 02;02.09 |
| 02;01.21 | 02;03.23 | 02;02.20 |
| 02;02.02 | 02;04.09 | 02;03.01 |
| 02;02.16 | 02;04.23 | 02;03.19 |
| 02;03.04 | 02;05.06 | 02;04.24 |
| 02;04.00 | 02;05.27 | 02;05.01 |
| 02;04.14 | 02;06.11 | 02;05.13 |
| 02;04.28 | 02;07.15 | 02;05.26 |
| 02;05.11 | 02;07.29 | — |
| 02;05.25 | 02;08.13 | — |
| 02;06.11 | 02;10.00 | — |
| 02;06.25 | — | — |

**TABLE III.** Summary of participants with atypical development. Data collected from children with DS (left), HI (center), and SLI (right). Age format is year;month.day. Within each column, participants are sorted by increasing age.

| Down's Syndrome (DS) | Hearing Impairment (HI) | Specific Language Impairment (SLI) |
| --- | --- | --- |
| Marcel 04;04.21 | Nita 04;05.12 | Rinanda 04;01.16 |
| John 08;01.27 | Berry 04;07.20 | Pierre 04;07.20 |
| Chantal 08;04.07 | Nanda 04;11.28 | Renzo 04;08.21 |
| Tineke 08;04.07 | Bjorn 05;02.27 | Monique 04;09.08 |
| Fiona 10;05.03 | Judith 05;06.18 | Wilma 05;01.02 |
| Sven 10;08.10 | Jenny 05;08.23 | Diana 05;01.04 |
| Sanne 13;00.02 | Wim 05;09.28 | Lieneke 05;03.07 |
| Antoinette 13;05.22 | Danielle 05;10.28 | Pascal 05;04.28 |
| Bert 13;08.17 | Jelle 05;11.19 | Mark 05;11.22 |
| Iris 15;01.09 | Sander 06;02.08 | Bertus 06;00.10 |
| Pieter 15;02.07 | Charlotte 06;03.20 | Pim 06;00.13 |
| Eline 15;06.16 | Jeroen 06;05.23 | Hessel 06;00.24 |
| Ronnie 16;03.16 | Sander 06;08.27 | Jelle 06;01.13 |
| Erik 16;07.17 | Mike 06;09.25 | Joep 06;01.26 |
| Anita 16;10.01 | Corine 06;11.17 | Ramon 06;02.10 |
| Martijn 17;01.09 | Merel 07;02.12 | Joost 06;07.22 |
| Michael 18;02.29 | Marco 07;09.13 | Pjotr 07;00.18 |
| Marius 18;05.02 | Kalinka 08;08.11 | Sjouke 07;01.26 |
| Lisa 18;07.18 | Peter 09;00.15 | Kees 07;04.19 |
| Katrijn 18;11.05 | Thomas 03;11.14 | Lennart 08;01.17 |

## Vocabulary size

*N*^*W*^ : A single graph, 𝒢, extracted from a single conversation session with one of the participants will contain as many nodes as unique words were uttered. The vocabulary size is the number of nodes in 𝒢:*N*^*W*^ ≡ |*V* |.

## Connected vocabulary fraction

*F*_*CC*_: During a session, a child might utter words that constitute a sentence of their own—e.g. an interjection, or an isolated ‘yes’. Such particles have no syntactic dependencies on any other words. Unless they are uttered within longer sentences in the same session, as we build the network they will appear as isolated nodes. Similarly, a sentence might contain words that are uttered only once throughout a whole session. This would result in an isolated graph, a tree, disconnected from all others. These are ways in which a syntax graph might possess several connected components. Let 𝒢_*CC*_ be the largest connected component of a graph. Let*V*_*CC*_ ⊂*V* be the number of nodes in the largest connected component. Then*F*_*CC*_ ≡ |*V*_*CC*_|*/*|*V*| is the fraction of the vocabulary that belongs to the largest connected component.

## Number of edges

*N*^*E*^: Note that, the way we constructed our graphs𝒢, we ignore repeated syntactic associations. The number of edges in 𝒢,*N*^*E*^ ≡ |*E*|, then measured the unique distinct syntax relationships instantiated during a conversation session.

## Diameter

*D*: Between any two words in a syntax network (say nodes*v*_*i*_,*v*_*j*_ ∈*V*) we can find the shortest path connecting them. Let us call*d*(*i, j*) to the distance of this path. The diameter of a network is the largest shortest distance between any two nodes, min_{*i,j*}_(*d*(*i, j*)). Paths between distant words are made up of chains of valid syntactic relationships. These, when pieced together, make up sentences. Often a network diameter gets smaller when the graph is dense and when there are several alternative ways to get from one node to another. In our context this would mean several valid chains of syntactic relationships—i.e. a dense and richly webbed route map of possible, grammatical sentences. This indicates a richness of syntactic combinations. Opposite to this, when a diameter is large, it means that some nodes can be reached through a unique and long path. This, for us, means that the corresponding words are just connected by a singular sentence, indicating somehow poor syntactic combinatorics.

## Degree

⟨*k*⟩ : The degree of a node,*k*_*i*_, is the number of other nodes to which it is connected,*k*_*i*_ = ∑ _*k*_*a*_*ij*_. In our context this refers to the number of unique syntax relationships that the corresponding word is involved in. To characterize the network at large, we are interested in the average degree, ⟨*k*⟩ ≡ ∑ _*i*_*k*_*i*_*/N*^*W*^. Thus, a larger average degree indicates that words are more flexible, involved in more syntactic relationships than in networks with a smaller average degree. This again indicates us the combinatorial richness of the syntax behind a network.

## Clustering

*C*: The clustering coefficient of a network is the fraction of complete triangles among all possible triangles, ACHTUNG!! Eq. here. A triangle is completed when two nodes (say*j* and*k*) that are neighbor of a third one (say*i*) are also connected to each other—hence the set*E* contains all three edges (*i, j*), (*j, k*), and (*i, k*). Clustering coefficient talks to us about local density of connections. In our context, a high clustering means that if a word is connected to other two, those two words also have a syntactic relationship in some context. Note that this can be rare in syntax. Take the sentences “I like chocolate” and “I like oranges”. The word ‘like’ is syntactically connected to both ‘chocolate’ and ‘oranges’, but it is unlikely that these two words will share a syntactic relationship in a naturally occurring sentence. The same reasoning applies for other syntactic relationships, which often involve words of different kinds (here a verb and a noun), while words of a same kind take more effort to combine. These reflections will be important when considering the coloring number of our graphs.

## Assortativity

*r*: Assortativity quantifies whether a node is similar to its neighbors regarding some property—usually degree. If, in a network, nodes with large degree tend to have neighbors who are also well connected, while vertices with few links are connected to each other, then that graph is assortative. If well connected nodes link to those with few connections, then the network is antiassortative. We use the Pearson correlation coefficient,*r*, between each node’s degree and that of its neighbors to measure assortativity [58]. Linguistically, since a word’s degree tells us about how many distinct syntactic connections that word can form, an assortative graph would mean that words with similar syntactic valence are linked together. That would be the case, e.g., if central connectors (such as pronouns or the verb to be) would preferentially link to each other. This is typically not the case in language networks. They are antiassortative, meaning that a few connector words tend to link together to nodes that have a lower capacity to combine with others.

## *k*-cores

A graph’s*k*-core is found by iteratively removing all nodes with degree less than*k* until no more nodes can be removed [59, 60]. Take 2-cores as an example: From a network, remove all nodes that have degree less than 2. In the resulting graph, some nodes might now have degree 1. We remove these next. We repeat, until no node is left with less than 2 neighbors. If no node is left at the end of this process, then the network does not have a 2-core (a typical syntax tree often does not have a 2-core). We find that most of our syntax graphs have got 2- and 3-cores of varied sizes, but few have 4- or higher*k*-cores. We call*K*_*k*_ to the set of nodes in a*k*-core and take the sizes |*K*_2_| and |*K*_3_| of 2- and 3-cores to characterize each network. We also take:

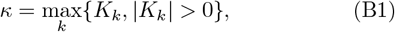

i.e. the*k* of the largest*k*-core.*k*-cores tell us whether a graph contains densely connected regions, as we can think of them as core sets of nodes that survive once peripheries are removed. Linguistically, the presence of a*k*-core would be telling us that a graph contains a dense, varied subset of syntactic relationships that are recurrent—i.e. that involve some same nodes repeatedly and in different situations.

## Cycles

Characterizing cycles is particularly challenging because they grow combinatorially as soon as networks show recurrence—i.e. as loops are completed. We need to introduce some technicalities to account for cycles properly, but the definitions below just formalize the intuition that a cycle is a circuit of nodes within the network that return to a same point. Syntactically, to have cycles present in a graph, we need that some same words have been used in different sentences with different combinations of words. To provide an example: “Mary loves the clouds” and “Some clouds rained on Mary” would close a loop involving 6 different words: “Mary”, “loves”, “the”, “clouds”, “rained”, “on”, “Mary”. Having a large number of cycles in a graph says that a high linguistic combinatorics is at play. Cycles might be longer or shorter. Long cycles in syntax networks reflect long linear constructions (i.e. richer phrases) that are connected by some words at their extremes. Those connectors need to be reused in different phrases (such as “Mary” and “clouds” above). We wanted to capture two properties of cycles: their abundance and lengths. To this end we need to introduce some notation: Given a network 𝒢 with its sets*V* and*E* of nodes and edges respectively, a cycle of this network is an ordered collection of vertices,*c* = {*v*(1), …,*v*(*n*)} such that*v*(*i*) ∈*V* for all*i* = 1, …, *n* and such that (*v*(*i*),*v*(*i* + 1)) ∈*E* for all*i* = 1, …, *n* ™ 1 and (*v*(*n*),*v*(1)) ∈*E*. (Note that*v*(*i*) labels vertices within the cycle while we used*v*_*i*_ to label vertives within*V* .) The cycle length is given by*n*. Let us call*E*(*c*) = {(*v*(*i*),*v*(*i* + 1)),*i* = 1, …, *n*}∪ {(*v*(*n*),*v*(1))} to the set of edges involved in cycle*c*. We define the addition of two cycles as the operation resulting from the exclusive or (xor) of the corresponding sets of edges. This is, given two cycles*c*_1_ and*c*_2_ with edges*E*(*c*_1_) and*E*(*c*_2_), we say that*c*_3_ =*c*_1_ +*c*_2_ with*E*(*c*_3_) =*E*(*c*_1_) ⊕*E*(*c*_2_). Nodes within*c*_3_ might be ordered differently to those in*c*_1_ and*c*_2_. With this, we say that a set,*C* ≡ {*c*_*l*_,*l* = 1, …, |*C*|}, of cycles in G forms a basis of its cycles if any cycle in 𝒢 can be generated by linear combinations (i.e. by summing) cycles in*C*. We refer to |*C*| as the basis size. If*n*_*j*_ is the size of the*j*-th cycle in*C*, then:

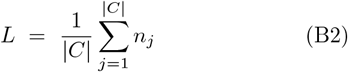

is the average length of cycles in*C*. Among all basis, some of them are minimal, meaning that they involve the least number of cycles that can generate all others. But due to their combinatorial explosion, finding minimal basis in a short time is prohibitive. Because of this, we evaluate cycle properties stochastically. Using a greedy algorithm [61] (implemented by Python’s networkx library), we generate 100 cycle basis of each network. We take the mean of their sizes and the mean of the average length of their cycles to enter our analysis.

## Constraints

We took three graph measurements that might be interpreted as finding maximal constraints on a network: the*maximal matching size*, the*maximal independent set*, and the*coloring number*. This was inspired by earlier studies on syntax networks which appreciated the importance of the coloring number [42].

## Maximal matching size

A matching of a graph is a set of edges without common vertices. The maximal matching size is the size of the largest matching. This property will be larger in graphs that present long linear chains or sparse trees, while a star o a clique will have maximal matching sizes of 1. Thus we expect that this property decays as a network becomes more small world. We compute this quantity as the fraction of edges with respect to the total.

## Maximal independent set

This property is the converse of the previous one—with nodes instead of edges. An independent set if a set of nodes without common vertices. The maximal independent set is the largest independent set. Contrary to the previous one, this property will be largest in a star and also large in a linear chain or a tree; but it can still be small in most small world graphs. This set is found with a stochastic greedy algorithm, so we run it 100 times. We take the quantity as the fraction of nodes in the independent set with respect to the total.

## Coloring number

This quantity represents the smallest number of colors that you would need to color the nodes of a graph such that two contiguous nodes are colored differently. This quantity is smallest for linear chains or stars, while maximal for a clique. Thus, all three constraints properties taken together, can explore across a range of complex graphs in which extremes we find cliques, stars, or linear chains.

## Jaccard similarity

The Jaccard similarity,*s*_*J*_, between two nodes tells us if their neighborhoods are similar [62]. In a syntax network, this quantity will be large between words that combine with similar words—e.g. those in a same grammatical class. Let*V*_*i*_ ⊂*V*, be the neighborhood of node*i*:*V*_*i*_ ≡ {*v*_*k*_ ∈*V*, (*v*_*i*_,*v*_*k*_) ∈*E*}. Then:

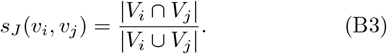

## Jaccard entropy

The property above averages Jaccard similarity over all nodes. There might be some nodes that tend to have neighborhoods that are more similar to those of others (think words that can link to any other), while others are less promiscuous in the kind of syntactic relationships they form. We wanted to capture a possible heterogeneity within the network. Since the Jaccard similarity is always a positive number, we captured this possible heterogeneity by normalizing the Jaccard similarity of all nodes, thus each value giving us a fraction within [0, 1], and all of them adding to 1. We then computed an entropy,*H*_*J*_. This entropy will be high if all nodes have similar Jaccard entropy, and smaller if there are some words with much higher values than others.

## Pagerank entropy

Similarly, we wanted to capture somehow a possible heterogeneity in the centrality of nodes within the network. To that end, we computed the eigenvector centrality of each node [31]. Since this is a positive number, we again normalized it and computed an entropy,*H*_*E*_.

## Appendix C: Supporting plots

**SUP. FIG. 1.**
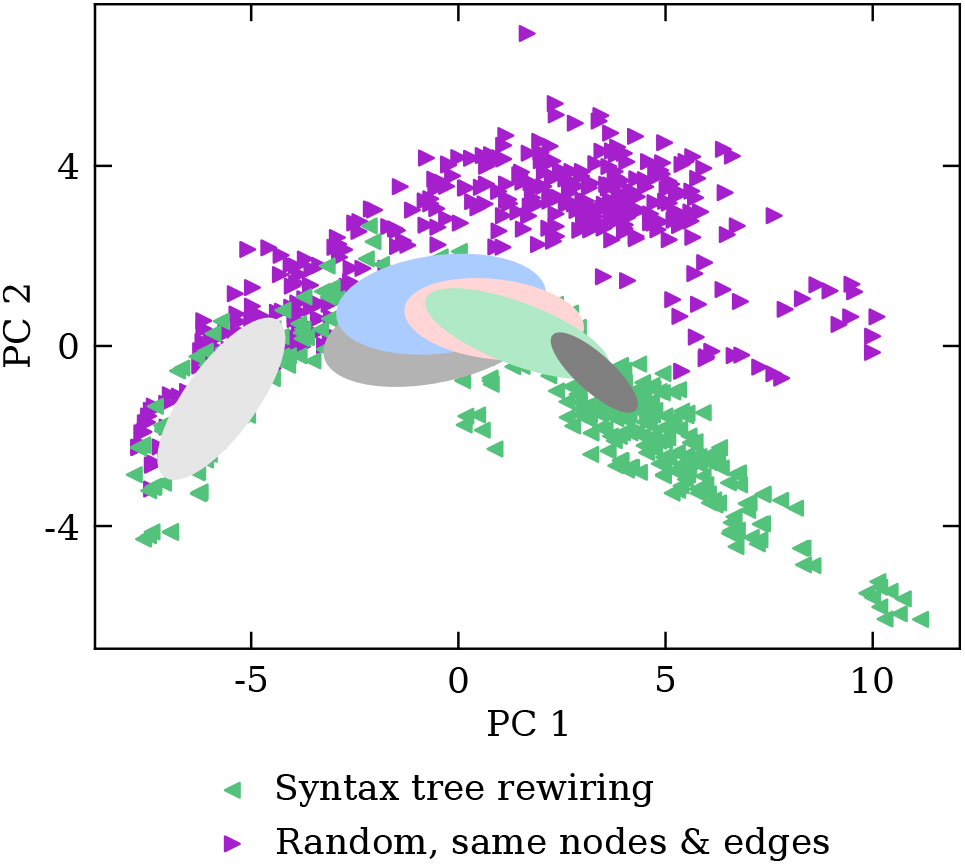
Syntax language morphospace with null-models. Marker colors and ellipsoids for empirical data is as in Fig. 4**a**. Longitudinal studies have been omitted for clarity. Null models are presented here to show how they stretch the morphospace, indicating that empirically-derived syntax networks are just a subset of all possible graphs. In other words, syntax networks constrain the kinds of connections between nodes that can exist. The rewiring (Rwr.) null model (leftward pointing triangles, darker green) falls fairly along the developmental trajectory. This null-mode is made by rewiring the list of edges of real syntax graphs, so we expect that it retains a fair amount of a syntax network structure—e.g. each word will have the same degree and be involved in as many syntactic relationships as the original. The random (Rmd.) null model (rightward pointing triangles, violet), instead, preserves much less of the original structure, as it consists of graphs with the same number of nodes and average degree. This null model explores a wider region of morphospace, meaning that it builds more different networks than those found empirically. Notably, vocabulary size is preserved with respect to the original. We see that these networks trace PC-1 well, but then diverge wildly alongside P-2, notably so after the peak of the developing trajectory. This suggests that the transition to the final developmental stage has associated some ability related to PC-2 (perhaps, e.g., to network diameter, notably with respect to connected vocabulary, Sup. Fig. 10) that random networks are unable to complete on their own.

**SUP. FIG. 2.**
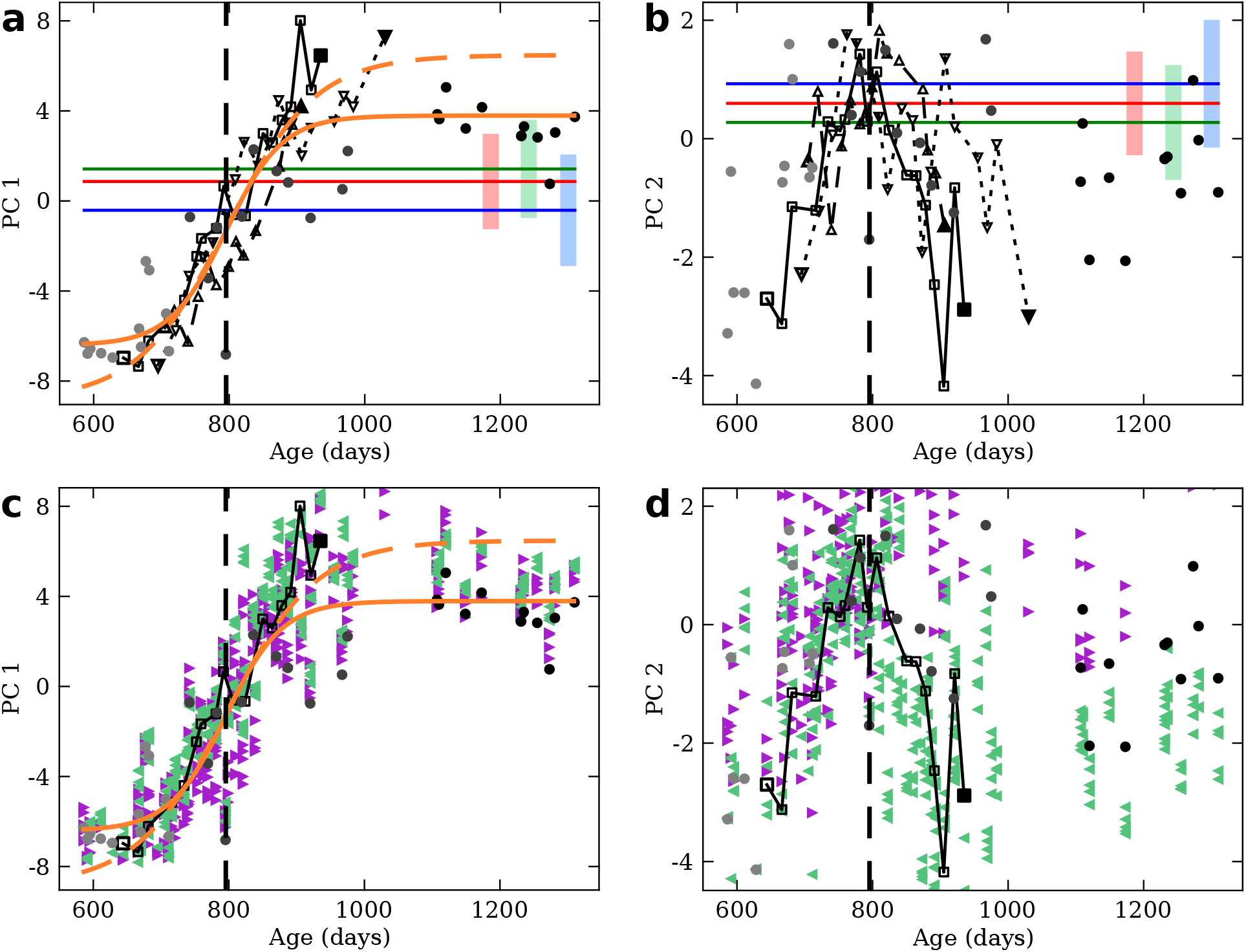
Principal components over time. Data points from typically developing children are shown as circles (squares or triangles for longitudinal studies) plotted at a specific age. Ages of typically developing children are well above the range of their typically developing counterparts. Also, the maturation level of atypically developing children does not necessarily correlate with age. Hence, for atypically developing children we show averages (solid horizontal bars) with standard deviations (light-colored rectangles). Color legends are as in Fig. 3. Null-model networks are plotted with the age of the network from which they were derived. Null-model colors are as in Sup. Fig. 1. A thick, dashed vertical line marks*t*_1*/*2_. **a** PC-1 over time for all empirical data collected. Orange curves show fits of Eq. 1 to data from all typically developing children (solid) or only longitudinal studies (dashed). **b** PC-2 over time for all empirical data collected. **c** PC-1 over time, alongside null-models. **d** PC-2 over time alongside null models.

**SUP. FIG. 3.**
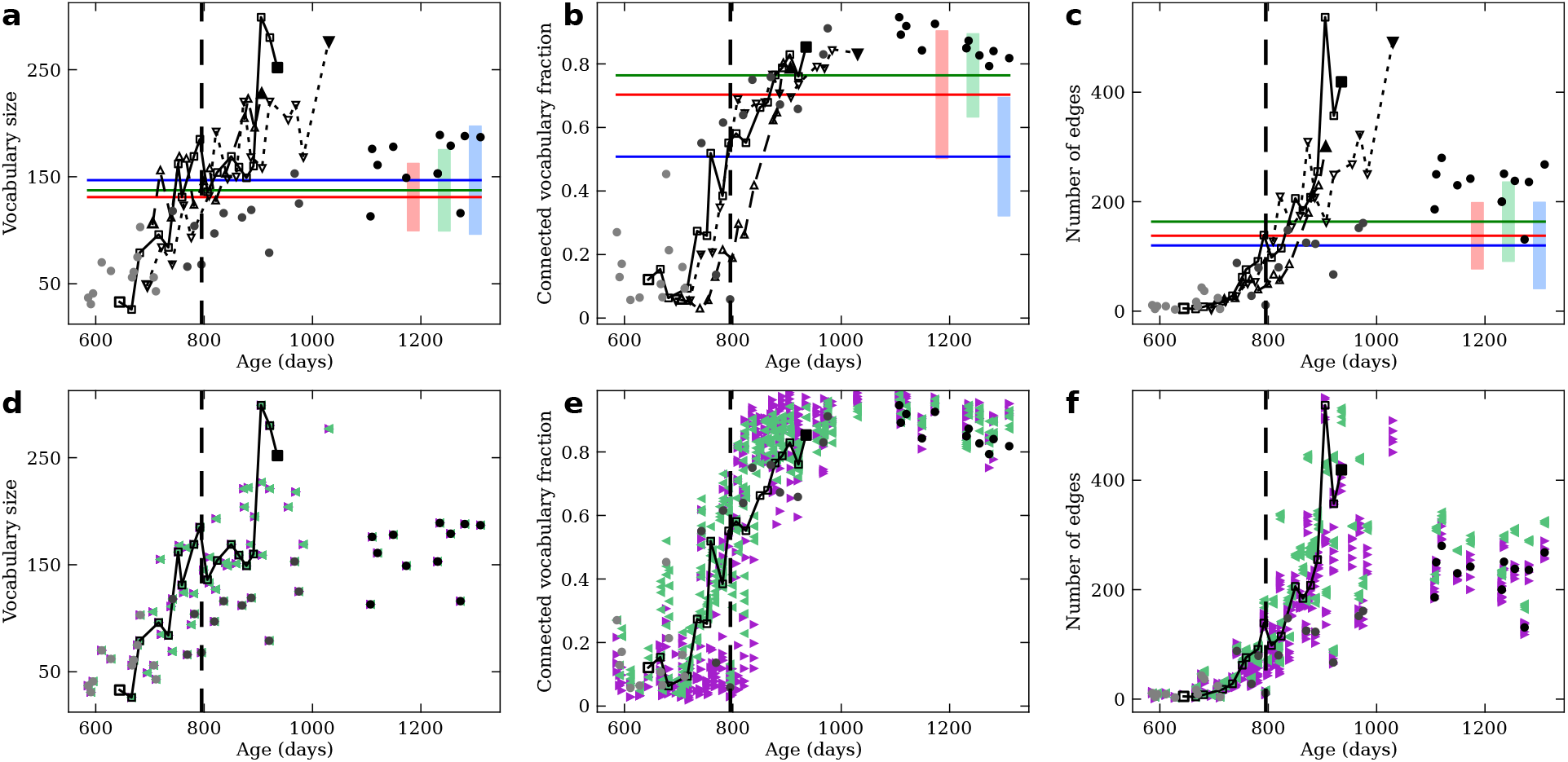
Properties related to network size over time. Data points from typically developing children are shown as circles (squares or triangles for longitudinal studies) plotted at a specific age. Ages of typically developing children are well above the range of their typically developing counterparts. Also, the maturation level of atypically developing children does not necessarily correlate with age. Hence, for atypically developing children we show averages (solid horizontal bars) with standard deviations (light-colored rectangles). Color legends are as in Fig. 3. Null-model networks are plotted with the age of the network from which they were derived. Null-model colors are as in Sup. Fig. 1. A thick, dashed vertical line marks*t*_1*/*2_. **a** Total vocabulary size over time for all empirical data collected. **b** Fraction of vocabulary that belongs to the network largest connected component over time. **c** Number of edges in the network over time. **d-f** Same plots, including null-model networks.

**SUP. FIG. 4.**
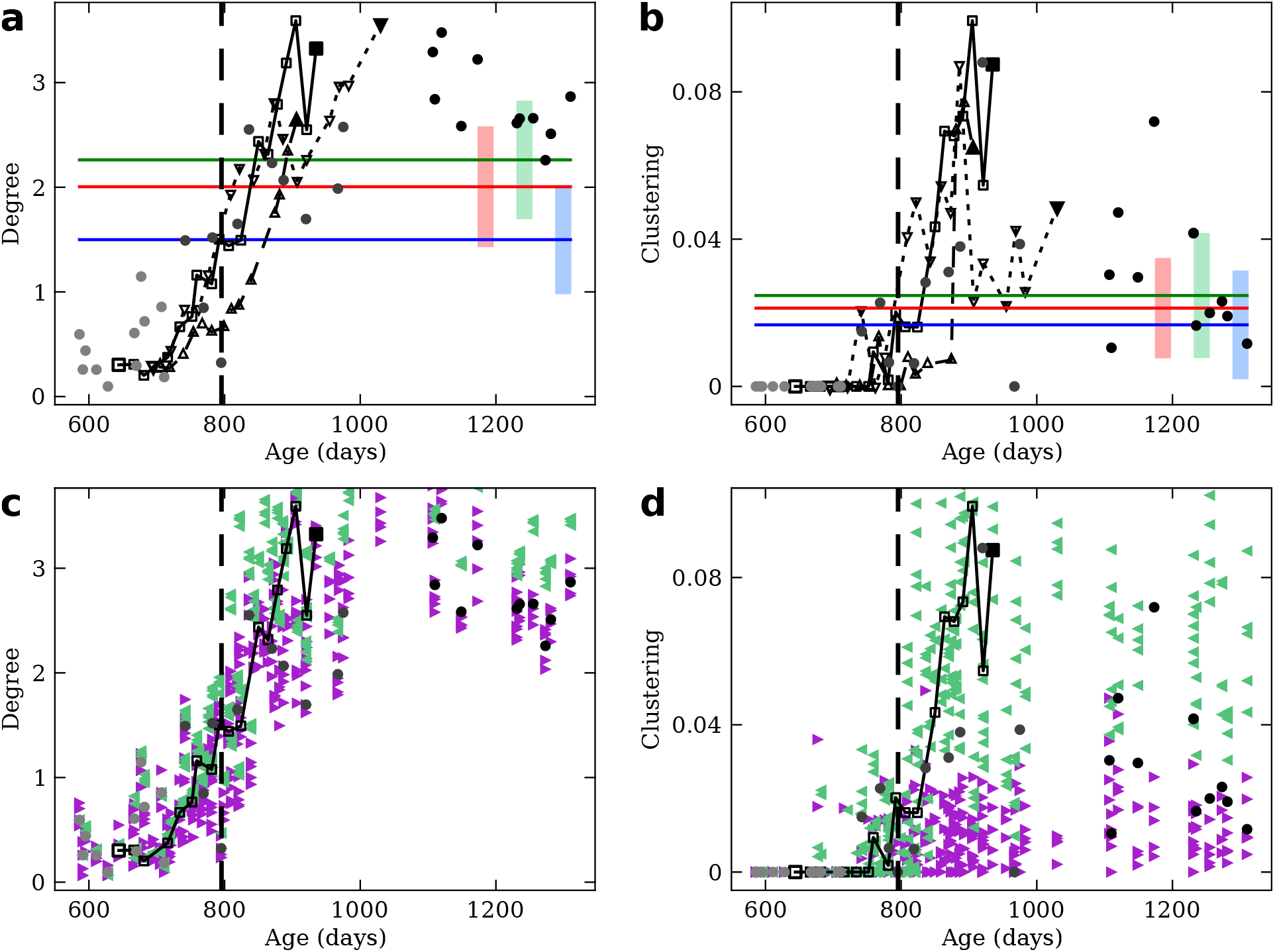
Degree and clustering over time. Data points from typically developing children are shown as circles (squares or triangles for longitudinal studies) plotted at a specific age. Ages of typically developing children are well above the range of their typically developing counterparts. Also, the maturation level of atypically developing children does not necessarily correlate with age. Hence, for atypically developing children we show averages (solid horizontal bars) with standard deviations (light-colored rectangles). Color legends are as in Fig. 3. Null-model networks are plotted with the age of the network from which they were derived. Null-model colors are as in Sup. Fig. 1. A thick, dashed vertical line marks*t*_1*/*2_. **a** Average node degree over time for all empirical data collected. **b** Average clustering coefficient over time. **c-d** Same plots, including nullmodel networks.

**SUP. FIG. 5.**
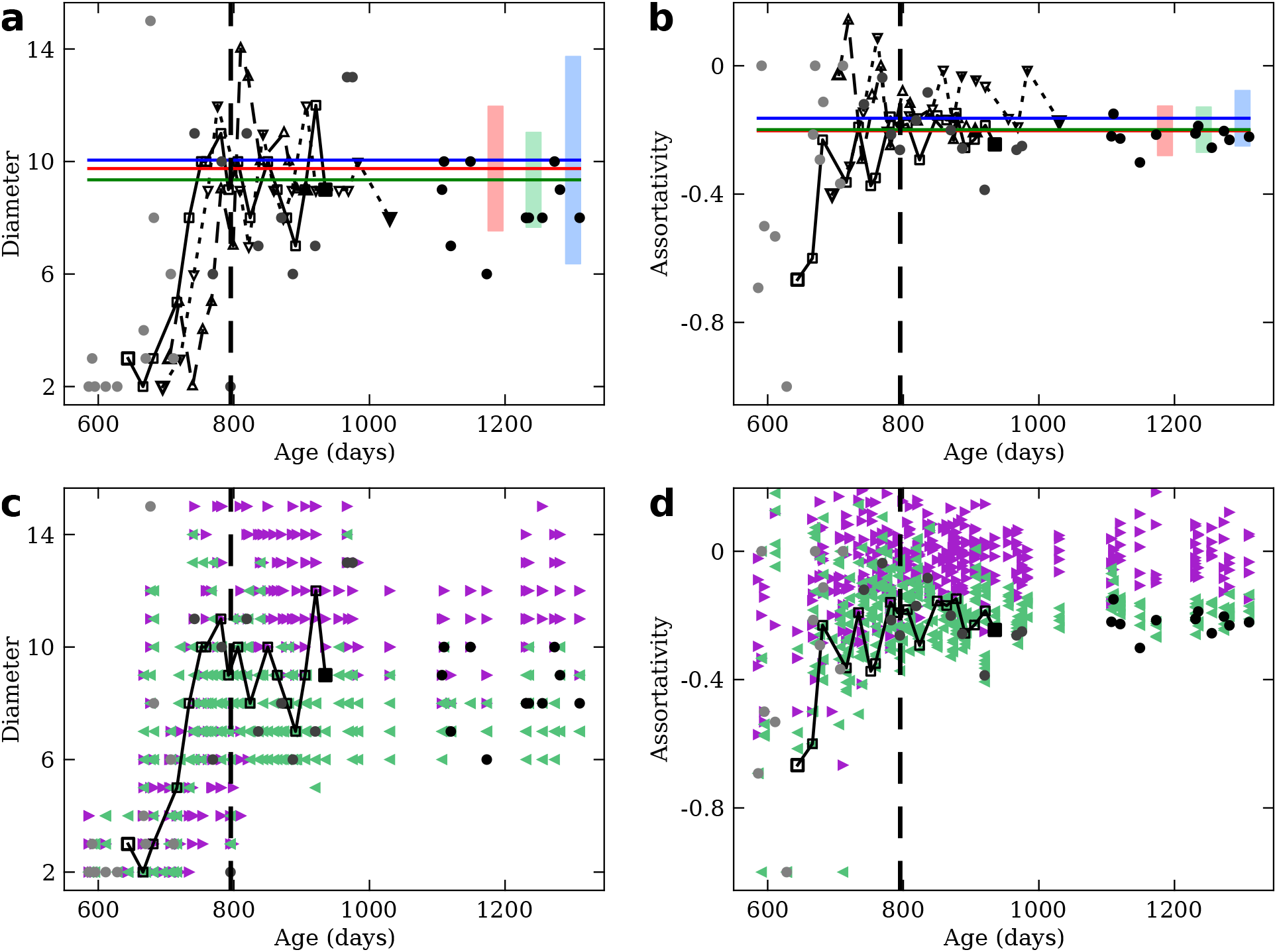
Diameter and assortativity over time. Data points from typically developing children are shown as circles (squares or triangles for longitudinal studies) plotted at a specific age. Ages of typically developing children are well above the range of their typically developing counterparts. Also, the maturation level of atypically developing children does not necessarily correlate with age. Hence, for atypically developing children we show averages (solid horizontal bars) with standard deviations (light-colored rectangles). Color legends are as in Fig. 3. Null-model networks are plotted with the age of the network from which they were derived. Null-model colors are as in Sup. Fig. 1. A thick, dashed vertical line marks*t*_1*/*2_. **a** Network diameter over time. **b** Assortativity over time. **c-d** Same plots, including null-model networks.

**SUP. FIG. 6.**
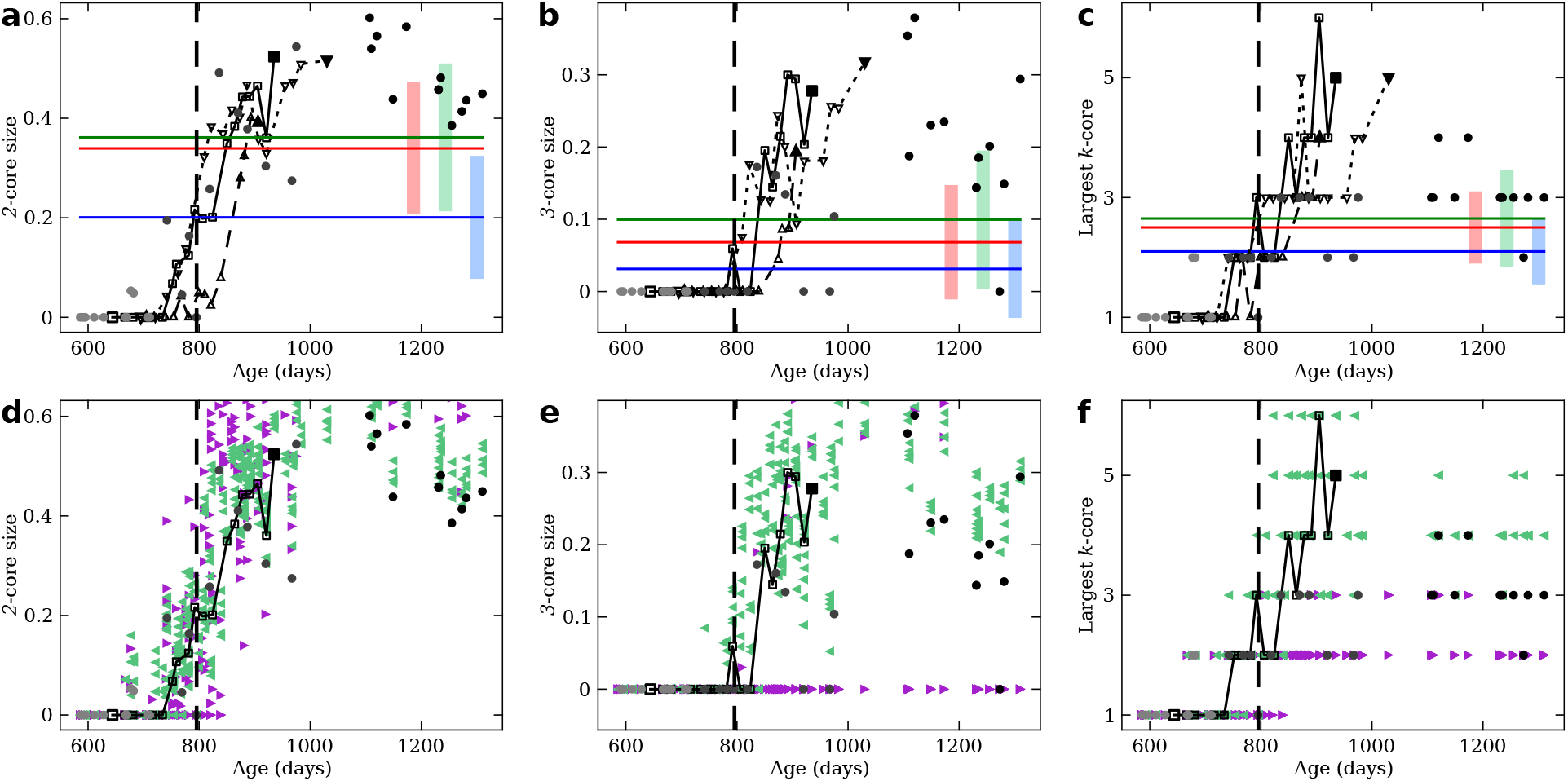
Measurements related to*k*-cores over time. Data points from typically developing children are shown as circles (squares or triangles for longitudinal studies) plotted at a specific age. Ages of typically developing children are well above the range of their typically developing counterparts. Also, the maturation level of atypically developing children does not necessarily correlate with age. Hence, for atypically developing children we show averages (solid horizontal bars) with standard deviations (light-colored rectangles). Color legends are as in Fig. 3. Null-model networks are plotted with the age of the network from which they were derived. Null-model colors are as in Sup. Fig. 1. A thick, dashed vertical line marks*t*_1*/*2_. **a** Size of the 2-core over time. **b** Size of the 3-core over time. **c** Largest non-empty*k*-core in the network, over time. **d-f** Same plots, including null-model networks.

**SUP. FIG. 7.**
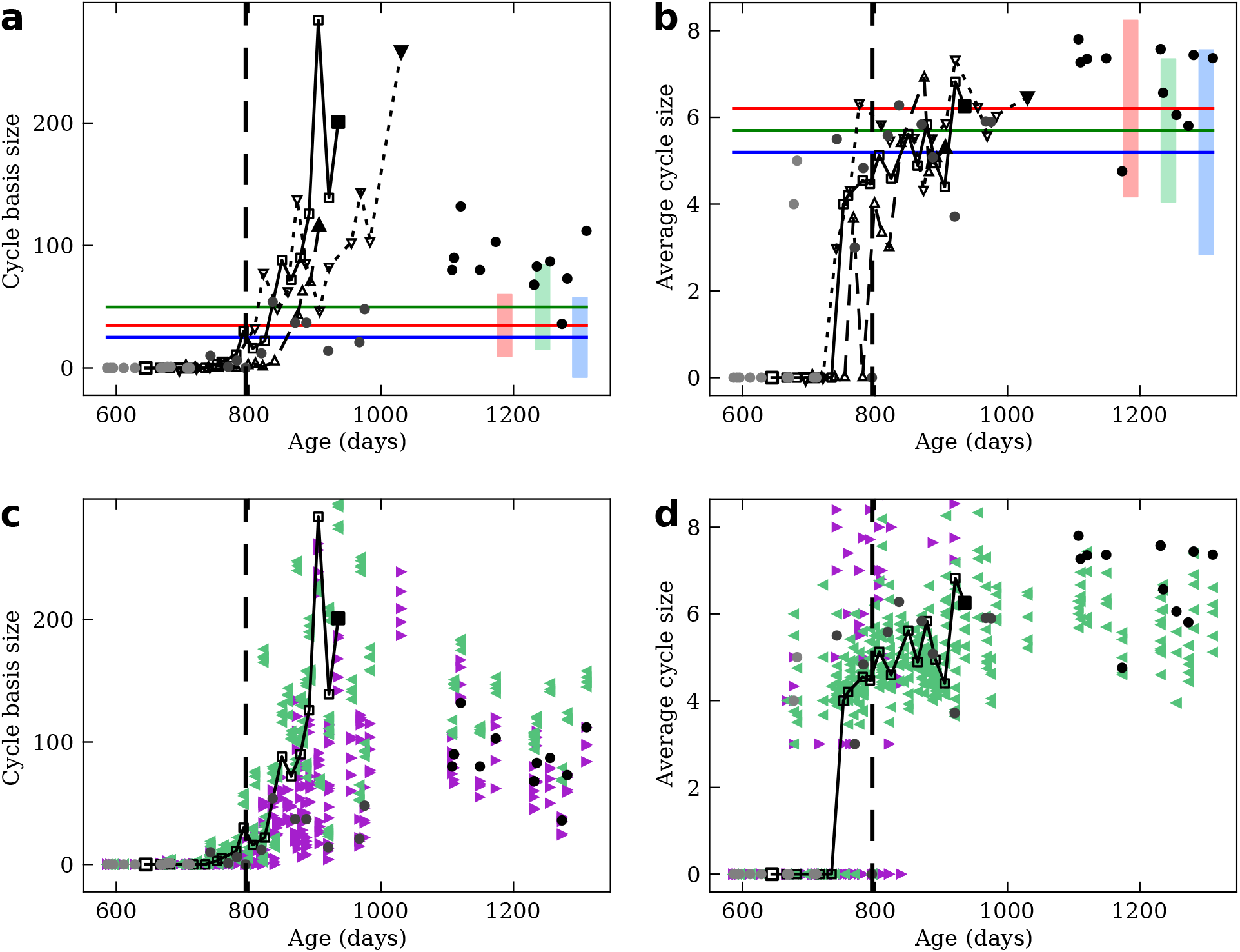
Measurements related to cycles over time. Data points from typically developing children are shown as circles (squares or triangles for longitudinal studies) plotted at a specific age. Ages of typically developing children are well above the range of their typically developing counterparts. Also, the maturation level of atypically developing children does not necessarily correlate with age. Hence, for atypically developing children we show averages (solid horizontal bars) with standard deviations (light-colored rectangles). Color legends are as in Fig. 3. Null-model networks are plotted with the age of the network from which they were derived. Null-model colors are as in Sup. Fig. 1. A thick, dashed vertical line marks*t*_1*/*2_. **a** Average size of cycle bases over time for all empirical data collected. **b** Average length of cycles in cycle bases over time. **c-d** Same plots, including null-model networks.

**SUP. FIG. 8.**
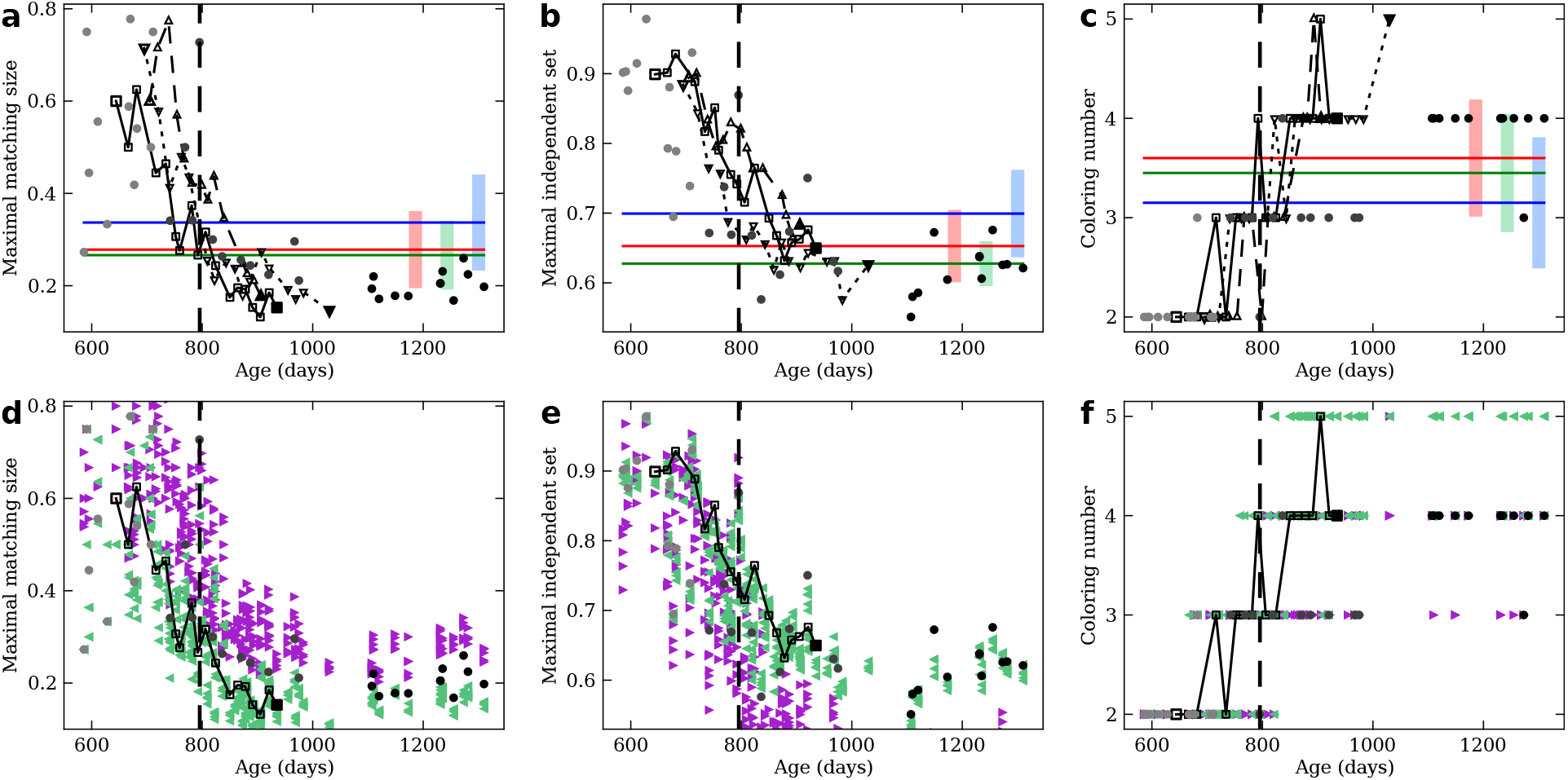
Measurements related to constraints imposed on graphs over time. Data points from typically developing children are shown as circles (squares or triangles for longitudinal studies) plotted at a specific age. Ages of typically developing children are well above the range of their typically developing counterparts. Also, the maturation level of atypically developing children does not necessarily correlate with age. Hence, for atypically developing children we show averages (solid horizontal bars) with standard deviations (light-colored rectangles). Color legends are as in Fig. 3. Null-model networks are plotted with the age of the network from which they were derived. Null-model colors are as in Sup. Fig. 1. A thick, dashed vertical line marks*t*_1*/*2_. **a** Maximal matching size over time for all empirical data collected. **b** Maximal independent set over time. **c** Coloring number over time. **d-f** Same plots, including null-model networks.

**SUP. FIG. 9.**
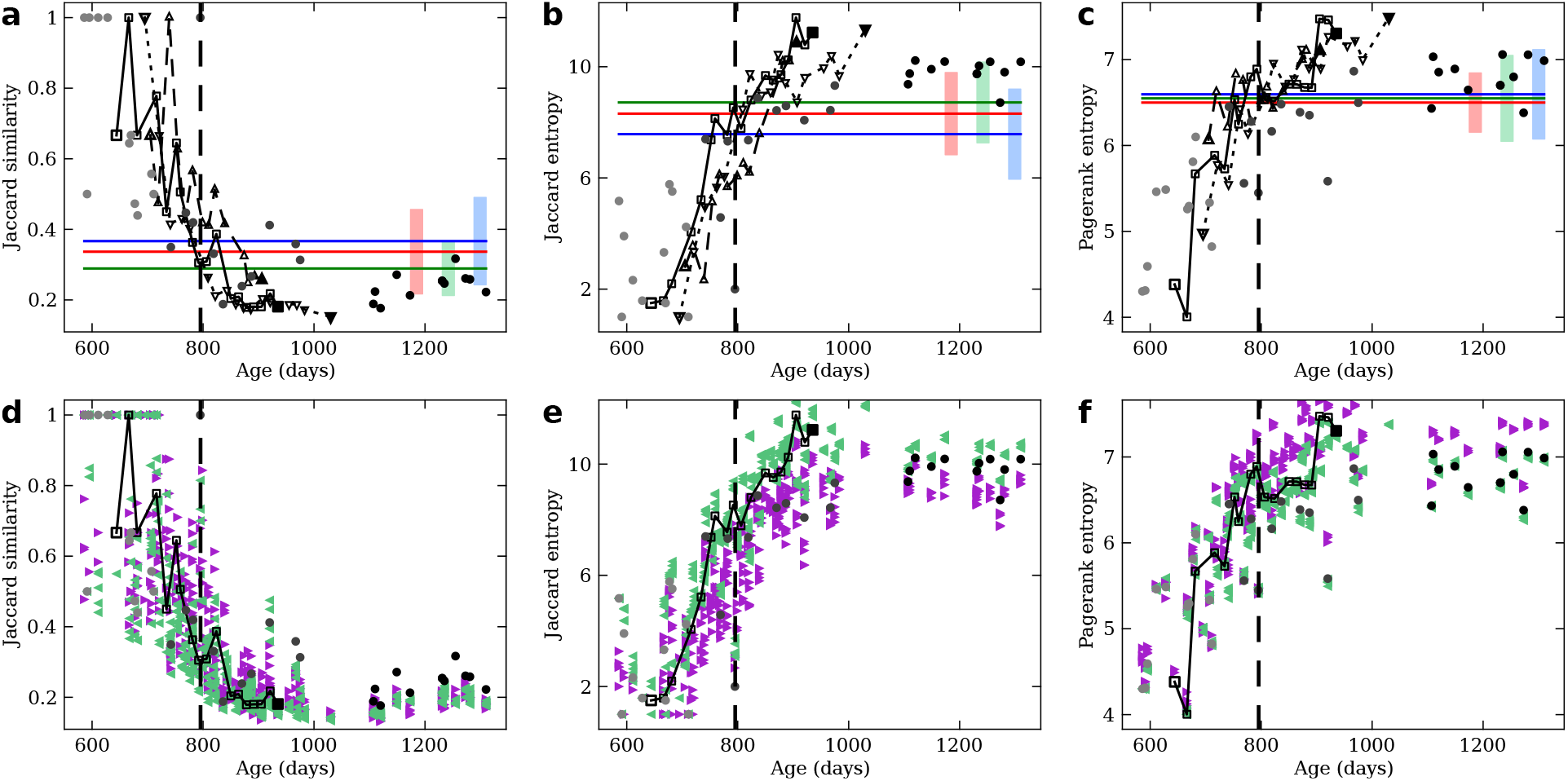
Measurements related to similarity between nodes over time. Data points from typically developing children are shown as circles (squares or triangles for longitudinal studies) plotted at a specific age. Ages of typically developing children are well above the range of their typically developing counterparts. Also, the maturation level of atypically developing children does not necessarily correlate with age. Hence, for atypically developing children we show averages (solid horizontal bars) with standard deviations (light-colored rectangles). Color legends are as in Fig. 3. Null-model networks are plotted with the age of the network from which they were derived. Null-model colors are as in Sup. Fig. 1. A thick, dashed vertical line marks*t*_1*/*2_. **a** Jaccard similarity over time for all empirical data collected. **b** Jaccard entropy over time. An increased Jaccard entropy means that nodes in a graph have more similar values of Jaccard similarity to each-other. Since panel **a** shows Jaccard similarity dropping, an increased Jaccard entropy means that nodes tend to be equally dissimilar to each-other. **c** Pagerank entropy over time. As with Jaccard entropy, this quantity raising means that eigenvector centrality of most nodes becomes more similar over time. In other words, differences in eigenvector centrality are larger for syntax networks in earlier developmental stages. **d-f** Same plots, including null-model networks.

**SUP. FIG. 10.**
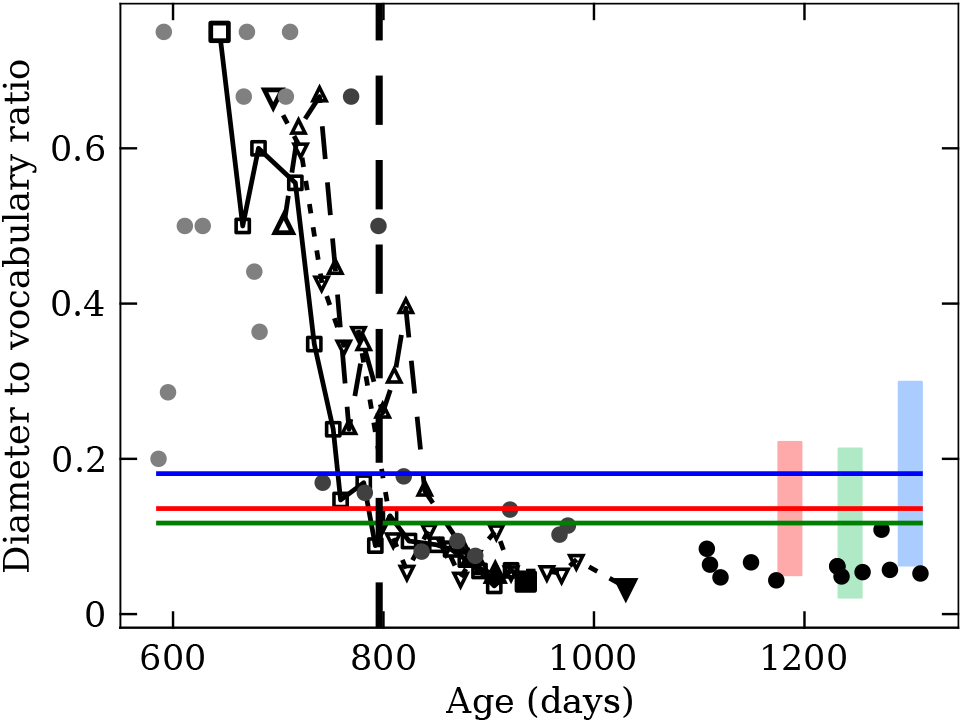
Ratio of network diameter to connected vocabulary over time. Data points from typically developing children are shown as circles (squares or triangles for longitudinal studies) plotted at a specific age. Ages of typically developing children are well above the range of their typically developing counterparts. Also, the maturation level of atypically developing children does not necessarily correlate with age. Hence, for atypically developing children we show averages (solid horizontal bars) with standard deviations (light-colored rectangles). Color legends are as in Fig. 3. A thick, dashed vertical line marks*t*_1*/*2_.

**SUP. FIG. 11.**
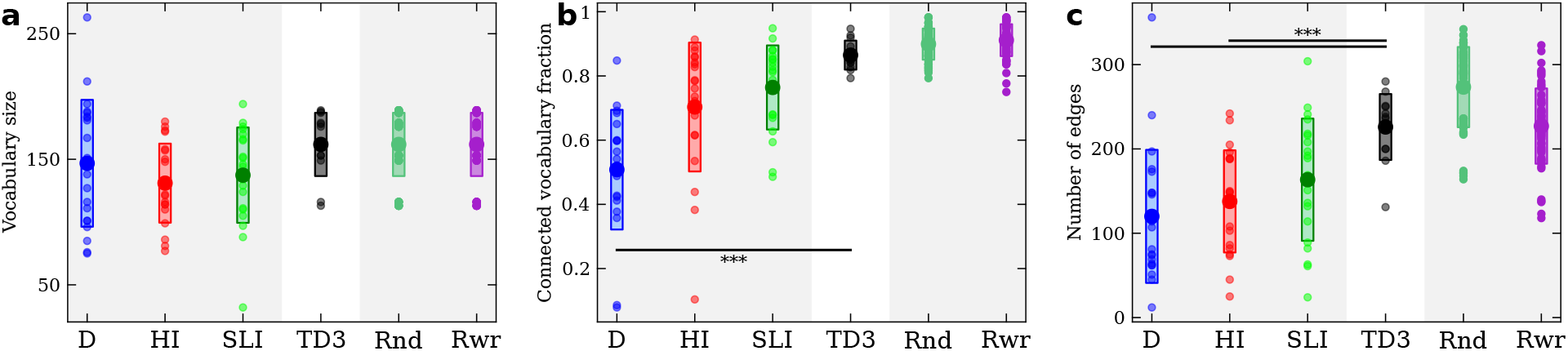
Distribution, averages, standard deviations, and significance of measurements of size for atypically developing children and null-models compared to oldest typically developing children. Each panel presents, for each group of networks, all empirically measured values of a property (small markers) together with their average (large markers), and standard deviation (light-colored boxes). Groups of networks presented are, from left to right: children with Down syndrome (DS, blue), hearing impairment (HI, red), specific language impairment (SLI, green), oldest cohort of typically developing children (TD3, black), Rnd. bootstrapped networks (darker green), and Rwr. bootstrapped network (violet). Significant differences are marked in comparisons between any group and TD3 with solid horizontal lines and three asterisks, ∗ ∗ ∗, indicating a *p*-value lower than 10^−3^. **a** Comparison of vocabulary sizes across groups. **b** Comparison of fractions of vocabulary in the largest connected component across groups. **c** Comparison of number of edges across groups.

**SUP. FIG. 12.**
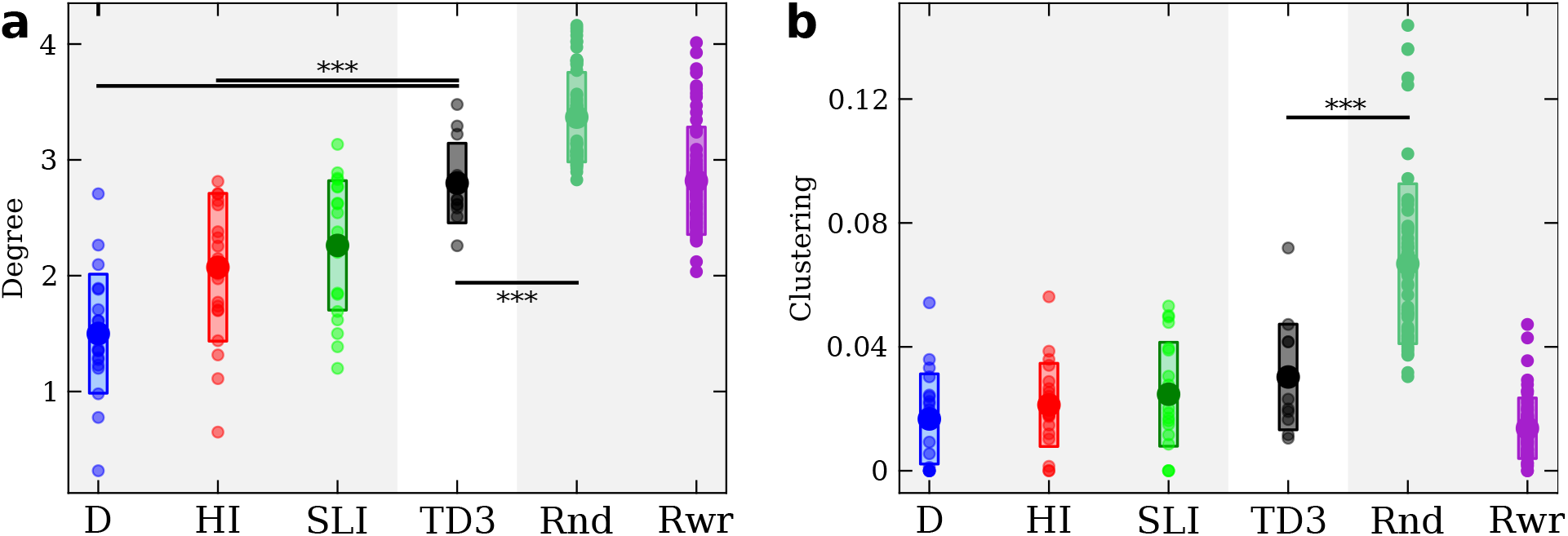
Distribution, averages, standard deviations, and significance of degree and clustering for atypically developing children and null-models compared to oldest typically developing children. Each panel presents, for each group of networks, all empirically measured values of a property (small markers) together with their average (large markers), and standard deviation (light-colored boxes). Groups of networks presented are, from left to right: children with Down syndrome (DS, blue), hearing impairment (HI, red), specific language impairment (SLI, green), oldest cohort of typically developing children (TD3, black), Rnd. bootstrapped networks (darker green), and Rwr. bootstrapped network (violet). Significant differences are marked in comparisons between any group and TD3 with solid horizontal lines and three asterisks, ∗ ∗ ∗, indicating a *p*-value lower than 10^−3^. **a** Comparison of degree across groups. **b** Comparison of clustering coefficient across groups.

**SUP. FIG. 13.**
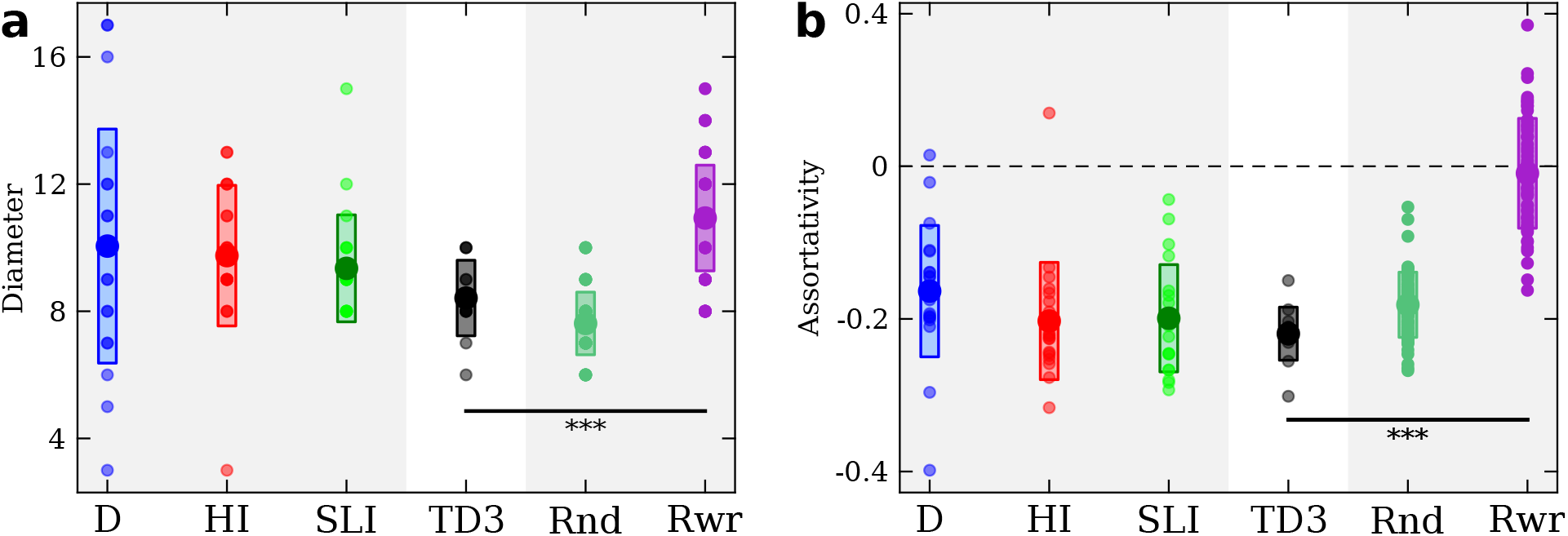
Distribution, averages, standard deviations, and significance of diameter and assortativity for atypically developing children and null-models compared to oldest typically developing children. Each panel presents, for each group of networks, all empirically measured values of a property (small markers) together with their average (large markers), and standard deviation (light-colored boxes). Groups of networks presented are, from left to right: children with Down syndrome (DS, blue), hearing impairment (HI, red), specific language impairment (SLI, green), oldest cohort of typically developing children (TD3, black), Rnd. bootstrapped networks (darker green), and Rwr. bootstrapped network (violet). Significant differences are marked in comparisons between any group and TD3 with solid horizontal lines and three asterisks, ∗ ∗ ∗, indicating a *p*-value lower than 10^−3^. **a** Comparison of diameter across groups. **b** Comparison of assortativity across groups.

**SUP. FIG. 14.**
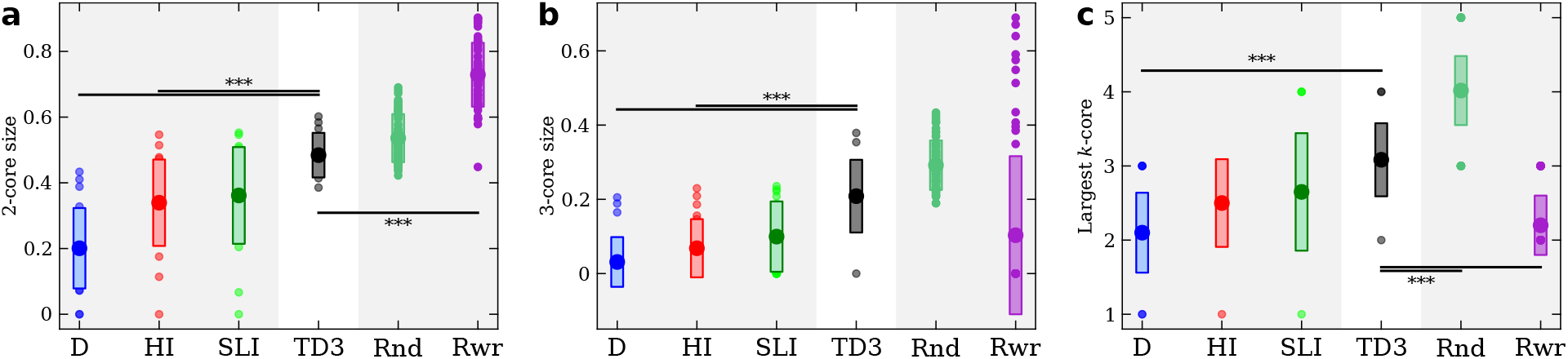
Distribution, averages, standard deviations, and significance of measurements related to*k*-cores for atypically developing children and null-models compared to oldest typically developing children. Each panel presents, for each group of networks, all empirically measured values of a property (small markers) together with their average (large markers), and standard deviation (light-colored boxes). Groups of networks presented are, from left to right: children with Down syndrome (DS, blue), hearing impairment (HI, red), specific language impairment (SLI, green), oldest cohort of typically developing children (TD3, black), Rnd. bootstrapped networks (darker green), and Rwr. bootstrapped network (violet). Significant differences are marked in comparisons between any group and TD3 with solid horizontal lines and three asterisks, ∗ ∗ ∗, indicating a *p*-value lower than 10^−3^. **a** Comparison of sizes of 2-cores across groups. **b** Comparison of sizes of 3-cores across groups. **c** Comparison of largest non-empty*k*-core across groups.

**SUP. FIG. 15.**
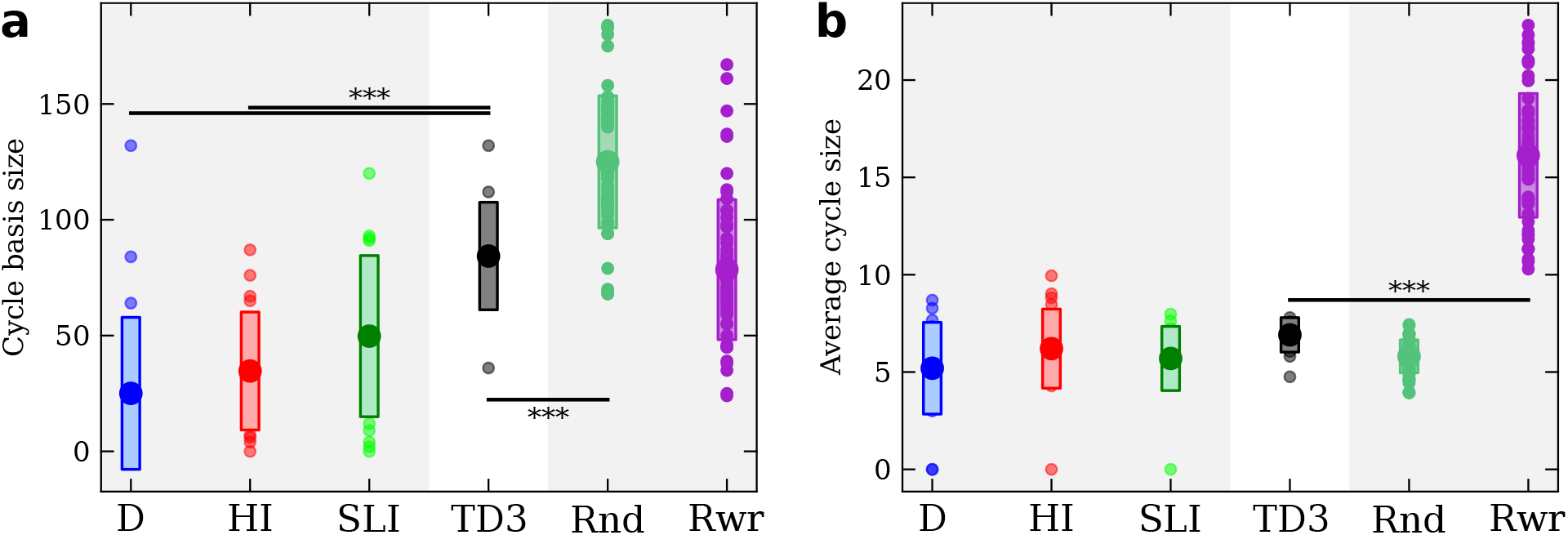
Distribution, averages, standard deviations, and significance of measurements related to cycles for atypically developing children and null-models compared to oldest typically developing children. Each panel presents, for each group of networks, all empirically measured values of a property (small markers) together with their average (large markers), and standard deviation (light-colored boxes). Groups of networks presented are, from left to right: children with Down syndrome (DS, blue), hearing impairment (HI, red), specific language impairment (SLI, green), oldest cohort of typically developing children (TD3, black), Rnd. bootstrapped networks (darker green), and Rwr. bootstrapped network (violet). Significant differences are marked in comparisons between any group and TD3 with solid horizontal lines and three asterisks, ∗ ∗ ∗, indicating a *p*-value lower than 10^−3^. **a** Comparison of average sizes of cycle bases across groups. **b** Comparison of average lengths of cycles in bases across groups.

**SUP. FIG. 16.**
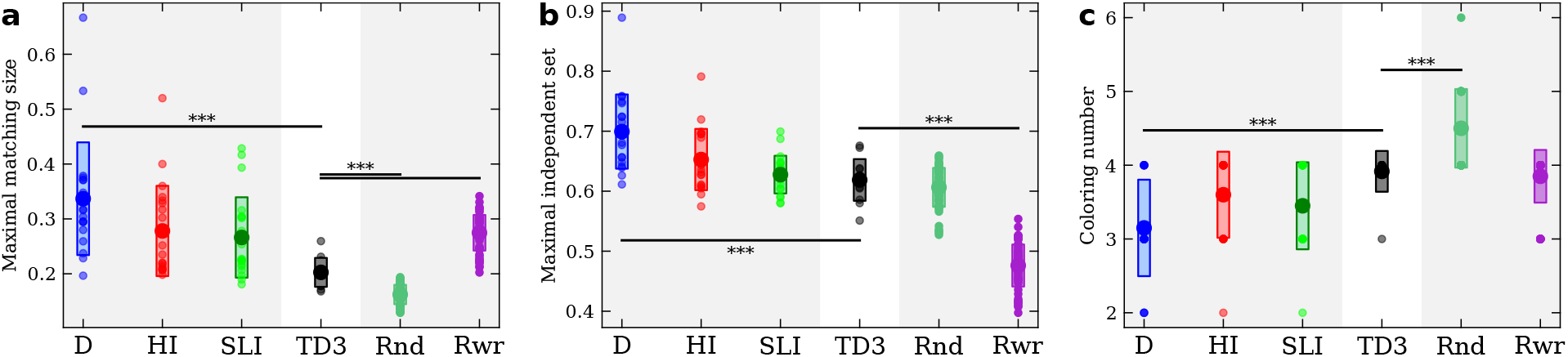
Distribution, averages, standard deviations, and significance of measurements related to constraints imposed on networks for atypically developing children and null-models compared to oldest typically developing children. Each panel presents, for each group of networks, all empirically measured values of a property (small markers) together with their average (large markers), and standard deviation (light-colored boxes). Groups of networks presented are, from left to right: children with Down syndrome (DS, blue), hearing impairment (HI, red), specific language impairment (SLI, green), oldest cohort of typically developing children (TD3, black), Rnd. bootstrapped networks (darker green), and Rwr. bootstrapped network (violet). Significant differences are marked in comparisons between any group and TD3 with solid horizontal lines and three asterisks, ∗ ∗ ∗, indicating a *p*-value lower than 10^−3^. **a** Comparison of maximal matching sizes across groups. **b** Comparison of maximal independent set sizes across groups. **c** Comparison of coloring numbers across groups.

**SUP. FIG. 17.**
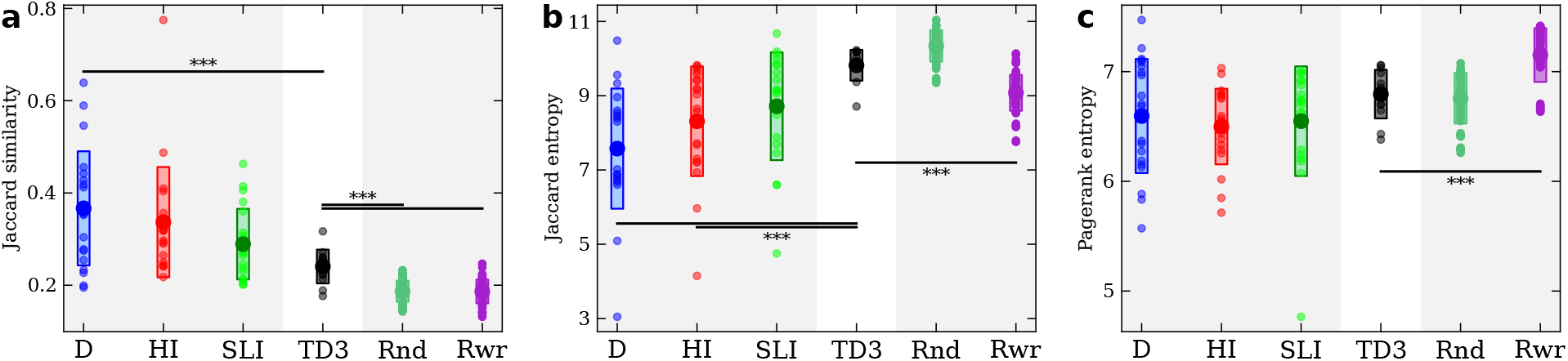
Distribution, averages, standard deviations, and significance of measurements of similarities between nodes for atypically developing children and null-models compared to oldest typically developing children. Each panel presents, for each group of networks, all empirically measured values of a property (small markers) together with their average (large markers), and standard deviation (light-colored boxes). Groups of networks presented are, from left to right: children with Down syndrome (DS, blue), hearing impairment (HI, red), specific language impairment (SLI, green), oldest cohort of typically developing children (TD3, black), Rnd. bootstrapped networks (darker green), and Rwr. bootstrapped network (violet). Significant differences are marked in comparisons between any group and TD3 with solid horizontal lines and three asterisks, ∗ ∗ ∗, indicating a *p*-value lower than 10^−3^. **a** Comparison of Jaccard similarity across groups. **b** Comparison of Jaccard entropy across groups. **c** Comparison of pagerank entropy across groups.

